# A squalene-hopene cyclase in *Schizosaccharomyces japonicus* represents a eukaryotic adaptation to sterol-independent anaerobic growth

**DOI:** 10.1101/2021.03.17.435848

**Authors:** Jonna Bouwknegt, Sanne J. Wiersma, Raúl A. Ortiz-Merino, Eline S. R. Doornenbal, Petrik Buitenhuis, Martin Giera, Christoph Müller, Jack T. Pronk

**Affiliations:** Department of Biotechnology, Delft University of Technology, Van der Maasweg 9, 2629 HZ Delft, The Netherlands; Center for Proteomics and Metabolomics, Leiden University Medical Center, 233 3ZA Leiden, The Netherlands; Department of Pharmacy, Center for Drug Research, Ludwig-Maximillians University Munich, Butenandtstraße 5-13, 81377 Munich, Germany

**Author notes:** Corresponding author: Jack T. Pronk, **Email:**. These authors have contributed equally to this work, and should be considered co-first authors (Jonna Bouwknegt, Sanne J. Wiersma). **Author Contributions:** JB, SJW and JTP designed experiments and wrote the draft manuscript. JB, SJW, RAOM, ESRD, PB, MG and CM performed experiments. JB, SJW, RAOM, MG and CM analyzed data. All authors have read and approved of the final manuscript. **Competing Interest Statement:** JB, SJW and JTP are co-inventors on a patent application that covers parts of this work. The authors declare no conflict of interest.

**Keywords:** sterols, yeast, anaerobic, hopanoids, *Schizosaccharomyces*

## Abstract

Biosynthesis of sterols, which are key constituents of canonical eukaryotic membranes, requires molecular oxygen. Anaerobic protists and deep-branching anaerobic fungi are the only eukaryotes in which a mechanism for sterol-independent growth has been elucidated. In these organisms, tetrahymanol, formed through oxygen-independent cyclization of squalene by a squalene-tetrahymanol cyclase, acts as a sterol surrogate. This study confirms an early report (Bulder (1971), Antonie Van Leeuwenhoek, 37, 353–358) that *Schizosaccharomyces japonicus* is exceptional among yeasts in growing anaerobically on synthetic media lacking sterols and unsaturated fatty acids. Mass spectrometry of lipid fractions of anaerobically grown *Sch. japonicus* showed the presence of hopanoids, a class of cyclic triterpenoids not previously detected in yeasts, including hop-22(29)-ene, hop-17(21)-ene, hop-21(22)-ene and hopan-22-ol. A putative gene in *Sch. japonicus* showed high similarity to bacterial squalene-hopene cyclase (SHC) genes and in particular to those of *Acetobacter* species. No orthologs of the putative *Sch. japonicus* SHC were found in other yeast species. Expression of the *Sch. japonicus* SHC gene (*Sjshc1*) in *Saccharomyces cerevisiae* enabled hopanoid synthesis and supported ergosterol-independent anaerobic growth, thus confirming that one or more of the hopanoids produced by SjShc1 can act as ergosterol surrogate in anaerobic yeast cultures. Use of hopanoids as sterol surrogates represents a previously unknown adaptation of eukaryotic cells to anaerobic growth. The fast sterol-independent anaerobic growth of *Sch. japonicus* is an interesting trait for developing robust fungal cell factories for application in anaerobic industrial processes.

**Significance statement:** Biosynthesis of sterols requires oxygen. This study identifies a previously unknown evolutionary adaptation in a eukaryote, which enables anaerobic growth in absence of exogenous sterols. A squalene-hopene cyclase, proposed to have been acquired by horizontal gene transfer from an acetic acid bacterium, is implicated in a unique ability of the yeast *Schizosaccharomyces japonicus* to synthesize hopanoids and grow in anaerobic, sterol-free media. Expression of this cyclase in S. *cerevisiae* confirmed that at least one of its hopanoid products acts as sterol-surrogate. The involvement of hopanoids in sterol-independent growth of this yeast provides new leads for research into the structure and function of eukaryotic membranes, and into the development of sterol-independent yeast cell factories for application in anaerobic processes.

## Introduction

Sterols are key constituents of canonical eukaryotic membranes, in which they influence integrity, permeability and fluidity^1,2^. The core pathway for sterol biosynthesis is highly conserved but the predominant final products differ for animals (cholesterol), plants (phytosterols) and fungi (ergosterol)^3^. Multiple reactions in sterol biosynthesis require molecular oxygen and no evidence for anaerobic sterol pathways has been found in living organisms or in the geological record^3^. The first oxygen-dependent conversion in sterol synthesis is the epoxidation of squalene to oxidosqualene by squalene monooxygenase. Oxidosqualene is subsequently cyclized to lanosterol, the first tetracyclic intermediate in sterol biosynthesis, in an oxygen-independent conversion catalysed by oxidosqualene cyclase (OSC). Molecular oxygen is also required for multiple subsequent demethylation and desaturation steps^4^. In fungi, synthesis of a single molecule of ergosterol from squalene requires 12 molecules of oxygen.

Deep-branching fungi belonging to the Neocallimastigomycota phylum are the only eukaryotes that have been unequivocally demonstrated to naturally exhibit sterol-independent growth under strictly anaerobic conditions. These anaerobic fungi contain a squalene-tetrahymanol cyclase (STC), which catalyzes oxygen-independent cyclization of squalene to tetrahymanol^5,6^. This pentacyclic triterpenoid acts as a sterol surrogate, and acquisition of a prokaryotic STC gene by horizontal gene transfer is considered a key evolutionary adaptation of Neocalllimastigomycetes to the strictly anaerobic conditions of the gut of large herbivores^7^. The reaction catalyzed by STC resembles oxygen-independent conversion of squalene to hopanol and/or other hopanoids by squalene-hopene cyclases (SHC)^8^, which are found in many prokaryotes^9,10^. Some prokaryotes synthesize tetrahymanol by ring expansion of hopanol, in a reaction catalyzed by tetrahymanol synthase (THS) for which the precise mechanism has not yet been resolved^11^.

Already in the 1950s, anaerobic growth of the industrial yeast and model eukaryote *Saccharomyces cerevisiae* was shown to strictly depend on sterol supplementation^12^ or use of intracellular stores of this anaerobic growth factor^13^. Similarly, fast anaerobic growth of *S. cerevisiae*, which is a key factor in its large-scale application in bioethanol production, wine fermentation and brewing^14,15^, requires availability of unsaturated fatty acids (UFAs). Biosynthesis of UFAs by yeasts requires an oxygen-dependent acyl-CoA desaturase^16^ and, in anaerobic laboratory studies, the sorbitan oleate ester Tween 80 is commonly used as UFA supplement^13,17^.

Per gram of yeast biomass, ergosterol and UFA synthesis require only small amounts of oxygen. Studies on these oxygen requirements therefore require extensive measures to prevent unintended oxygen entry into cultures. Even though most yeast species readily ferment sugars to ethanol under oxygen-limited conditions, only very few grow anaerobically on sterol- and UFA-supplemented media when such precautions are taken^18,19^. As opposed to Neocallimastigomycetes, no evolutionary adaptations to sterol-independent anaerobic growth have hitherto been reported for yeasts, or for ascomycete and basidiomycete fungi in general. We recently demonstrated that expression of an STC gene from the ciliate *Tetrahymena thermophila* supported tetrahymanol synthesis and sterol-independent growth of *S. cerevisiae*^20^. This result inspired us to re-examine a 1971 publication in which Bulder^21^ reported sterol- and UFA-independent growth of the dimorphic fission yeast *Schizosaccharomyces japonicus*. *Sch. japonicus* was originally isolated from fermented fruit juices^22,23^, and its potential for wine fermentation is being explored^24,25^. It shows marked genetic and physiological differences with other fission yeasts^26,27^ and has gained interest as a model for studying cell division dynamics and hyphal growth^28–30^. *Sch. japonicus* grows well at elevated temperatures and rapidly ferments glucose to ethanol^31^. A low sterol content, control of membrane fluidity via chain length of saturated fatty acids and respiratory deficiency may all reflect adaptations of *Sch. japonicus* to low-oxygen environments^21,31–33^. However, the report by Bulder^21^ stating that *Sch. japonicus* can grow anaerobically without sterol supplementation has not been confirmed or further investigated.

An ability of *Sch. japonicus* to grow in the absence of an exogenous supply of sterols would raise urgent questions on the molecular and evolutionary basis for this trait, which is extremely rare among eukaryotes. Despite the current absence of experimental evidence, it has been proposed that oxygen-independent sterol synthesis is theoretically possible^34^. Alternatively, sterol-independent growth of *Sch. japonicus* might depend on synthesis of an as yet unidentified sterol surrogate, or on membrane adaptations that involve neither sterols nor sterol surrogates. In addition to these fundamental scientific questions, independence of anaerobic growth factors is a relevant trait for large-scale industrial applications of yeasts, as exemplified by ‘stuck’ brewing fermentations caused by depletion of intracellular sterols and/or UFA reserves^35,36^.

The goals of the present study were to reinvestigate the reported ability of *Sch. japonicus* to grow anaerobically without sterol supplementation and to elucidate its molecular basis. In view of reported challenges in avoiding oxygen contamination in laboratory cultures of yeasts^19,20,37,38^, we first reassessed anaerobic growth and lipid composition of *Sch. japonicus* in the presence and absence of ergosterol. After identifying a candidate SHC gene in *Sch. japonicus*, we investigated its role in anaerobic growth by its expression in *S. cerevisiae*. In addition, we tested the hypothesis of Bulder^39^ that *Sch. japonicus* is able to synthesize UFAs in an oxygen-independent pathway.

## Results

### Anaerobic growth of *Sch. japonicus* without ergosterol and UFA supplementation

In anaerobic laboratory cultures of yeasts, biosynthetic oxygen requirements are easily obscured by unintended entry of minute amounts of oxygen^19,20,37,38^. Furthermore, upon transfer from aerobic to anaerobic conditions, some yeasts continue growing on media without sterols or UFAs by mobilizing intracellular reserves^20,37^. To check if such complications affected conclusions from an early literature report on sterol- and UFA-independent anaerobic growth of *Sch. japonicus*^21^, we performed serial-transfer experiments in an anaerobic chamber^37^ using phosphate-buffered synthetic medium with glucose as carbon source (SMPD), with and without supplementation of ergosterol and/or Tween 80 as source of UFAs.

To deplete reserves of sterols and/or UFAs in aerobically grown cells, anaerobic pre-cultures were grown on SMPD with an increased glucose concentration (50 g L^−1^), and lacking ergosterol and Tween 80. In these pre-cultures, growth of *S. cerevisiae* CEN.PK113-7D ceased after 4.8 doublings (Figure 1A), when less than half of the glucose had been consumed (SI Appendix, Table S1). Under the same conditions, *Sch. japonicus* CBS5679 completed 6.1 doublings (Figure 1B) and, while full glucose depletion was intentionally avoided to prevent excessive flocculation and sporulation, it had consumed almost 90 % of the glucose (SI Appendix, Table S1). Samples from the anaerobic pre-cultures were transferred to SMPD (20 g L^−1^ glucose) supplemented with different combinations of ergosterol and/or Tween 80. Consistent with earlier reports^37^, *S. cerevisiae* showed virtually no anaerobic growth on SMPD without ergosterol and Tween 80 (Figure 1A). In contrast, *Sch. japonicus* showed maximum specific growth rates of 0.26 to 0.30 h^−1^ and reached optical densities of 4 to 5 in all media tested (Figure 1B; SI Appendix, Table S2). This anaerobic growth was sustained upon two consecutive transfers in SMPD lacking either Tween 80, ergosterol, or both (Figure 1B). These results confirmed Bulder’s^21^ conclusion that *Sch. japonicus* can grow anaerobically in the absence of sterol and UFA supplementation. Remarkably, *Sch. japonicus* grew substantially slower in aerobic cultures (0.19 h^−1^; SI Appendix, Figure S1) than in anaerobic cultures (Figure 1B; SI Appendix, Table S2).

**Figure 1.**
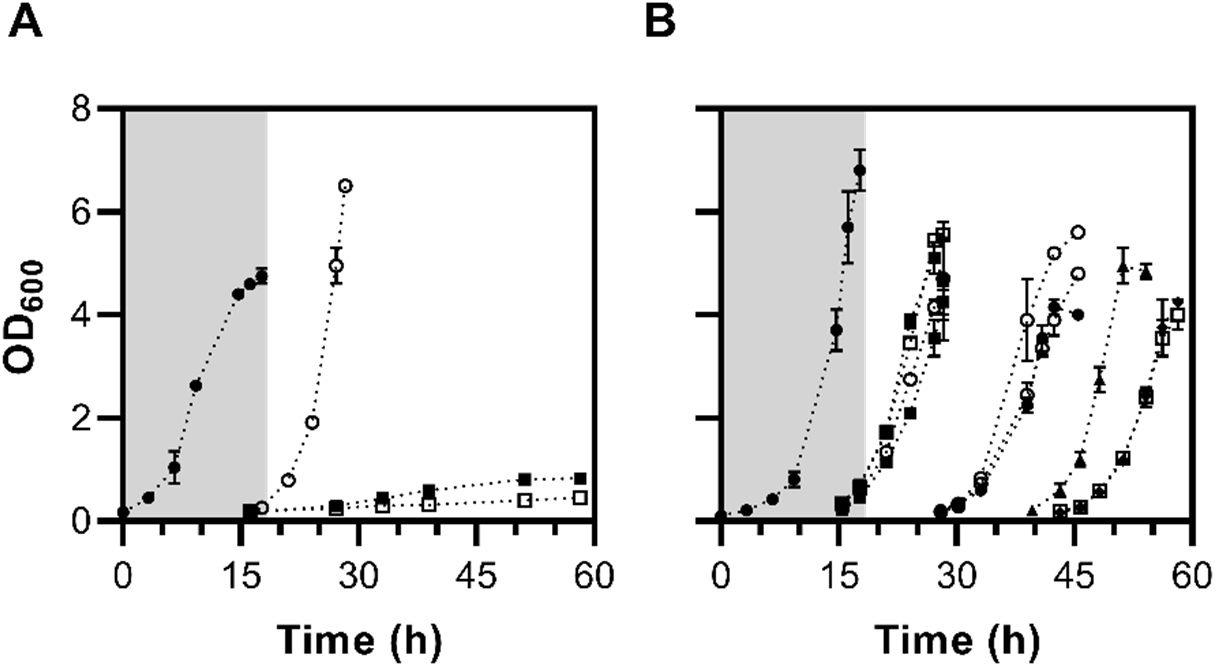
Anaerobic growth of *S. cerevisiae* CEN.PK113-7D and *Sch. japonicus* CBS5679 with different ergosterol and UFA (Tween 80) supplementation in a dark anaerobic chamber. Anaerobic pre-cultures on SMPD on 50 g L^−1^ glucose (closed circles, grey shading) were grown until the end of the exponential phase. (**A**) After the anaerobic pre-culture, *S. cerevisiae* was transferred to SMPD (20 g L^−1^ glucose) supplemented with either Tween 80 and ergosterol (open circles), Tween 80 only (closed squares) or neither Tween 80 nor ergosterol (open squares). (**B**) *Sch. japonicus* was grown on the same media as *S. cerevisiae* and additionally on medium containing ergosterol but not Tween 80 (closed triangles). *Sch. japonicus* cultures supplemented with only Tween 80, only ergosterol, and those without supplements were serially transferred to fresh media with the same composition in the anaerobic chamber. Data are represented as average ± SEM of measurements on independent duplicate cultures for each combination of yeast strain and medium composition.

### Absence of ergosterol and UFAs in anaerobically grown *Sch. japonicus*

To further investigate sterol- and UFA-independent anaerobic growth of *Sch. japonicus*, lipid fractions were isolated from anaerobic cultures and analyzed by gas chromatography with flame ionization detection (GC-FID). UFAs were detected in aerobically grown biomass, but not in anaerobic cultures grown on SMPD without Tween 80 supplementation (Figure 2; SI Dataset S01). These results showed that fast anaerobic growth of *Sch. japonicus* on UFA-free medium did not, as suggested by Bulder^39^, reflect oxygen-independent UFA synthesis. Total fatty-acid contents of aerobically and anaerobically grown biomass were similar, but anaerobically grown biomass showed higher contents of FA 10:0, FA 16:0 and FA 18:0 and lower contents of FA 26:0. In aerobically grown *Sch. japonicus* biomass, no FA 16:1 was detected and levels of FA 18:1 were higher than in Tween 80 supplemented anaerobic cultures (Figure 2).

**Figure 2.**
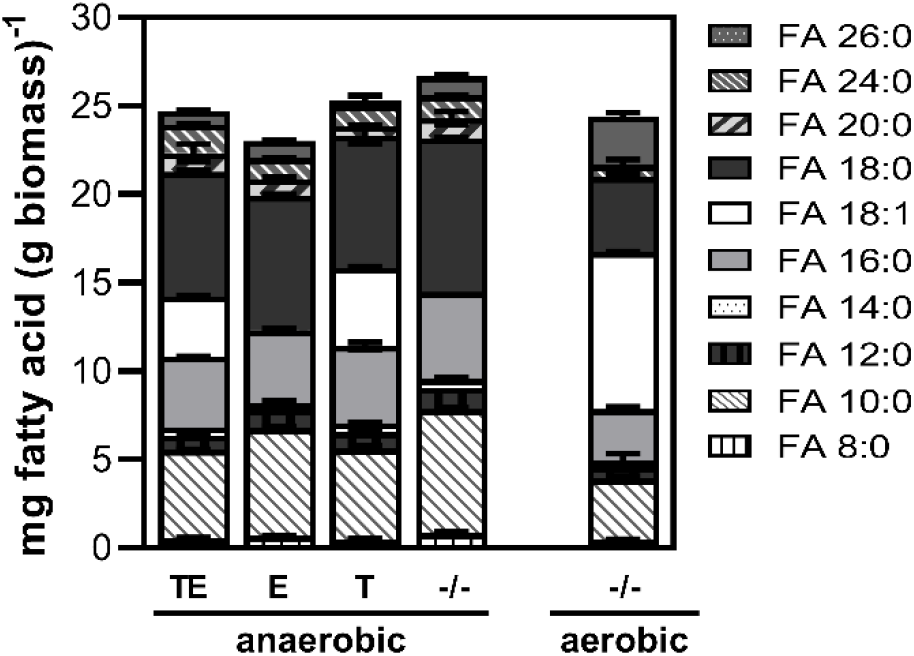
Quantification of fatty acids in *Sch. japonicus* CBS5679 biomass. *Sch. japonicus* CBS5679 was grown in SMPD with 20 g L^−1^ glucose. Under anaerobic conditions, cultures were supplemented with Tween 80 and ergosterol (TE), only ergosterol (E), only Tween 80 (T) or neither of those supplements (−/−). Data are shown for the first anaerobic culture following the anaerobic pre-culture. Aerobic cultures of *Sch. japonicus* were grown in SMPD without supplements (−/−). Data are represented as average ± SEM of measurements on independent duplicate cultures for each cultivation condition. Detailed information on data presented in this figure and additional anaerobic transfers are provided in SI Dataset S01.

*S. cerevisiae* biomass, grown anaerobically on SMPD with Tween 80 and ergosterol, was used as a reference for analysis of triterpenoid compounds and contained squalene, ergosterol and a small amount of lanosterol (Figure 3A). Similarly prepared triterpenoid fractions of anaerobic *Sch. japonicus* cultures that were supplemented with only Tween 80 did not contain detectable amounts of ergosterol or lanosterol. Instead, in addition to squalene, gas chromatography-mass spectrometry (GC-MS) revealed several compounds that were not observed in anaerobically grown *S. cerevisiae* (Figure 3B). We hypothesized that these compounds were as yet unidentified triterpenoids synthesized by *Sch. japonicus* that might serve as sterol surrogates. Tetrahymanol, which has not been found in wild-type yeasts but does occur in several other eukaryotes^6,40,41^, did not match any of the detected peaks based on its relative retention time (RRT, with cholestane as reference^20^).

**Figure 3.**
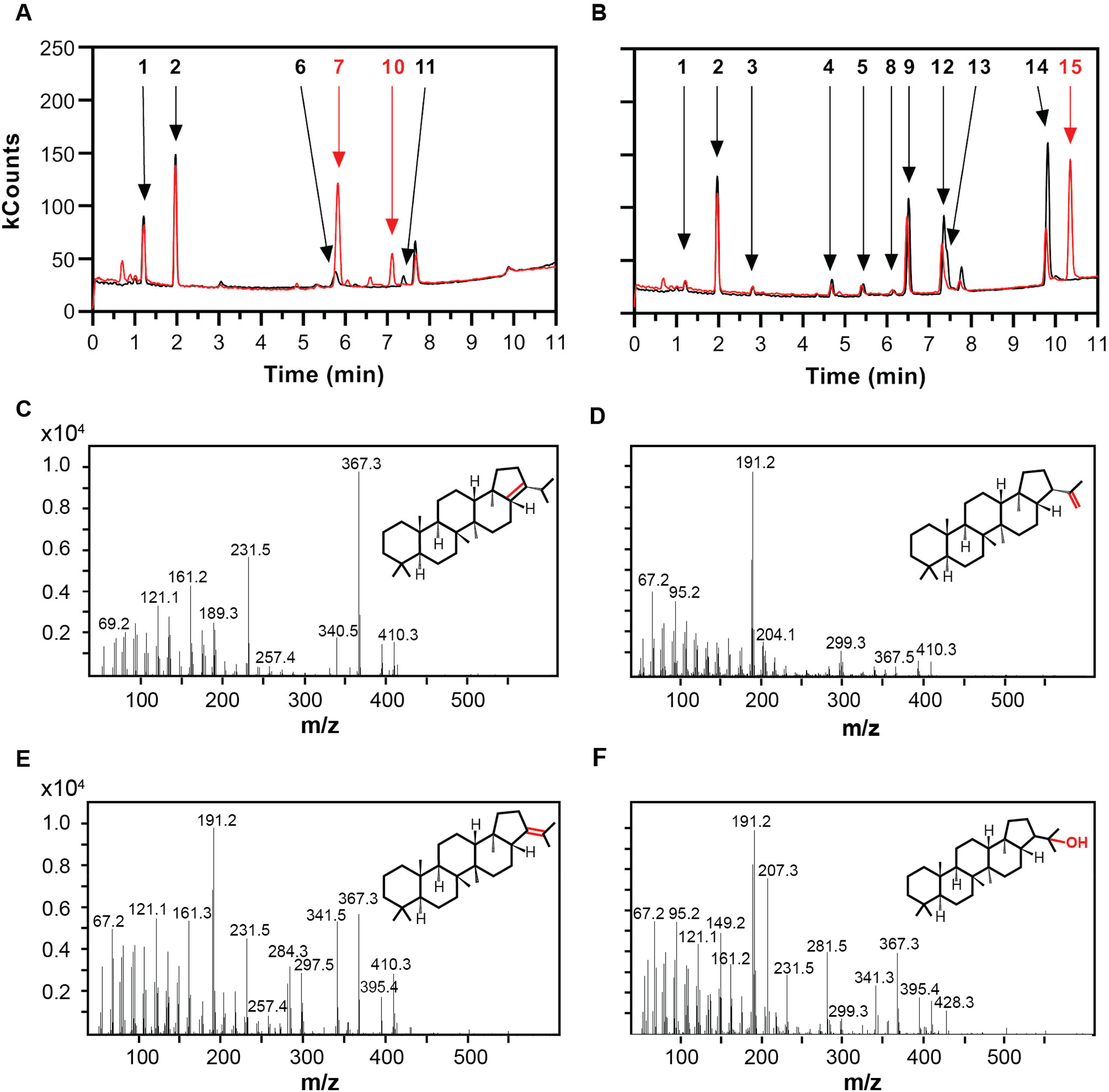
Gas chromatography-mass spectrometry (GC-MS) analysis of triterpenoid fractions of anaerobically grown yeast biomass. Anaerobic cultures were harvested in early stationary phase. Triterpenoids were extracted for GC-MS analysis and injected immediately (black lines) or after silylation (red lines). (**A**) *S. cerevisiae* CEN.PK113-7D grown anaerobically on medium supplemented with Tween 80 and ergosterol, (**B**) *Sch. japonicus* CBS5679 was grown on medium supplemented with only Tween 80. Numbers indicate the following compounds: **1**, squalene; **2**, 5α-cholestane (internal standard); **3**, squalene epoxide; **4**, hop-17(22)-ene; **5**, unidentified component; **6**, ergosterol; **7**, ergosterol-TMS-ether; **8**, unidentified component; **9**, unidentified component, possibly a tricyclic intermediate; **10**, lanosterol-TMS-ether; **11**, lanosterol; **12**, hop-22(29)-ene (diploptene); **13**, hop-21(22)-ene; **14**, hopan-22-ol (diplopterol); **15**, hopan-22-ol-TMS-ether. (**C-F**) Mass spectra and structures of identified hopanoid compounds in the triterpenoid fraction of anaerobically grown biomass of *Sch. japonicus*. (**C**) Hop-17(21)-ene, (**4**). (**D**) Hop-22(29)-ene, (**12**). (**E**) Hop-21(22)-ene, (**13**). (**F**) Hop-22-anol (diplopterol), (**14**).

### Predicted *Sch. japonicus* proteins resemble bacterial squalene-hopene cyclases

To identify potential sources of triterpenoids in *Sch. japonicus,* amino acid sequences of three characterized triterpene cyclases were used as queries to search the predicted proteomes of *Sch. japonicus* strains yFS275^26^ and CBS5679. Specifically, sequences of an oxidosqualene cyclase (OSC) from *Sch. pombe*^42^ (SpErg7; UniProt accession Q10231), a squalene-hopene cyclase (SHC) from *Acidocaldarius alicyclobacillus*^43^ (AaShc; P33247), and a squalene-tetrahymanol cyclase (STC) from *Tetrahymena thermophila*^5,20^ (TtThc1; Q24FB1), were used for a HMMER^44^ homology search. For each *Sch. japonicus* strain, this search yielded a sequence with significant homology to OSC, and another with significant homology to the SHC (Table 1). Consistent with the lipid analysis results, neither strain yielded a clear STC homolog.

**Table 1:** Homology search results using amino acid sequences of characterized triterpenoid cyclases against *Sch. japonicus* proteomes. Query coverage percentage and E-values were obtained with HMMER3^44^, identity percentages were calculated with Clustal^90^.

| Query | Accession of subject sequence | Query coverage (%) | E-value | Identity (%) |
| --- | --- | --- | --- | --- |
| *Subject proteome: Sch. japonicus* yFS275 |  |  |  |  |
| SpErg7<sup>a</sup> (Q10231) | B6JW54 | 99.7 | 0.0 | 65.9 |
| AaShc<sup>b</sup> (P33247) | B6K412 | 99.2 | $2.7 \times 10^{-141}$ | 38.1 |
| *Subject proteome: Sch. japonicus* CBS5679 |  |  |  |  |
| SpErg7<sup>a</sup> (Q10231) | SCHJC_A005630 (SjErg7) | 99.7 | 0.0 | 65.7 |
| AaShc<sup>b</sup> (P33247) | SCHJC_C003990 (SjShc1) | 98.9 | $2.5 \times 10^{-139}$ | 37.8 |

<sup>a</sup>Protein sequence of Erg7 of *Schizosaccharomyces pombe*
<sup>b</sup>Protein sequence of Shc of *Acidocaldarius alicyclobacillus*

To explore the phylogeny of the *Sch. japonicus* OSC and SHC homologs, SpErg7, AaSHC, and TtThc1 were used as queries for HMMER searches against all eukaryotic and all bacterial protein sequences in Universal Protein Resource (UniProt) reference proteomes (see SI Appendix, Table S3 for additional sequences obtained from TrEMBL). This search yielded 1480 eukaryotic and 348 bacterial cyclase homologs from a total number of 764 and 1826 species, respectively (SI Dataset S02). Sequences of cyclase homologs from 28 eukaryotic and 20 bacterial ‘species of interest’ were further analyzed. These species (Table S3) were either deep-branching anaerobic fungi^45^, Schizosaccharomycetes^26^ or species included in a previous publication on phylogeny of triterpene cyclases^46^. The selected cyclase homologs (SI Dataset S03) were subjected to multiple sequence alignment and used to generate a maximum-likelihood phylogenetic tree (Figure 4; SI Dataset S04).

**Figure 4.**
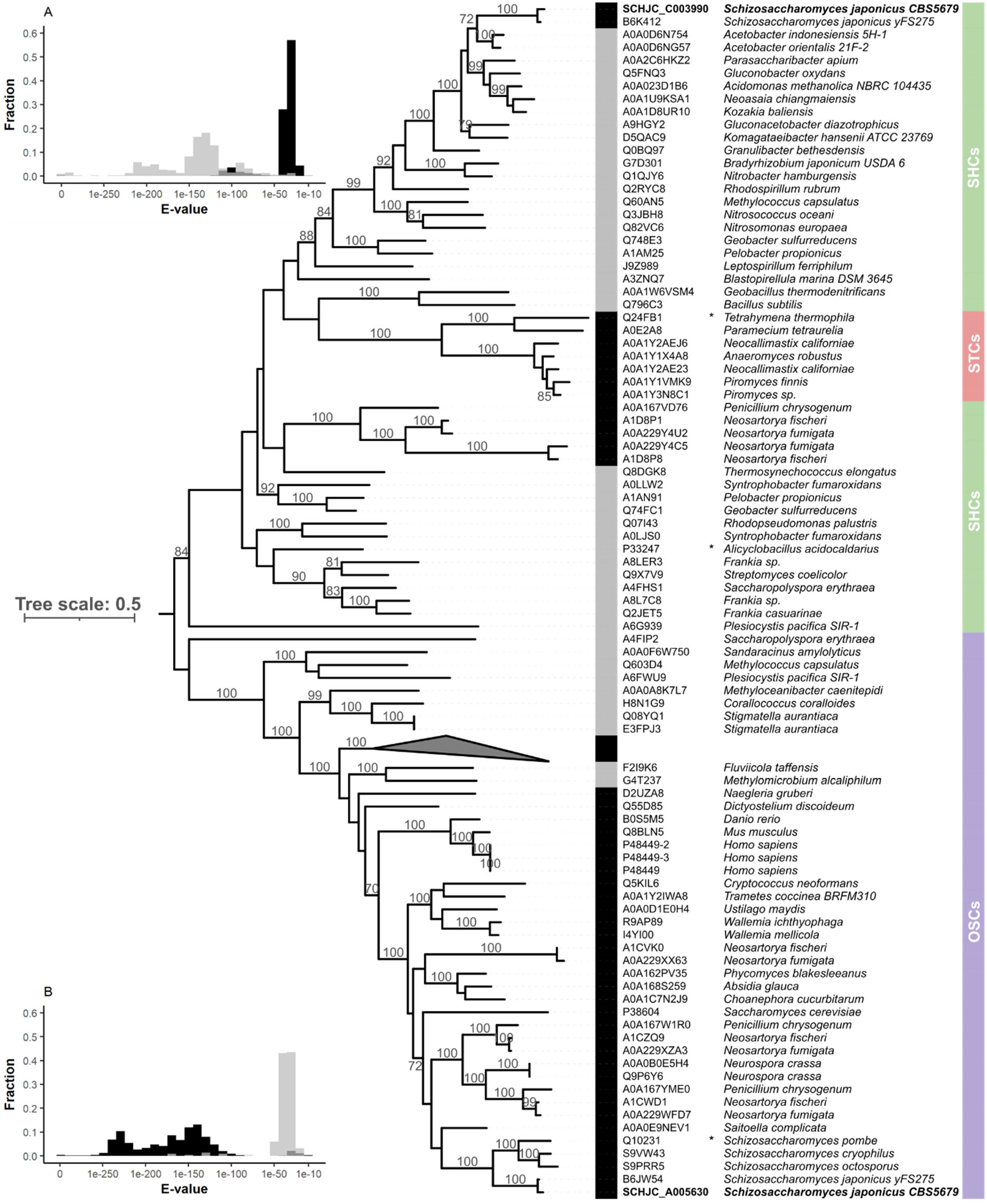
Maximum-likelihood phylogenetic tree of selected triterpenoid cyclases. The colored bar indicates different types of cyclases: green, squalene hopene cyclases (SHCs); red, squalene tetrahymanol cyclases (STCs); purple, and oxidosqualene cyclases (OSCs). Sequences were obtained from a systematic homology search using the characterised cyclases marked with an asterisk (*Acidocaldarius alicyclobacillus* AaShc, P33247, SHC; *Tetrahymena thermophila*, TtThc1, Q24FB1, STC; *Schizosaccharomyces pombe* SpErg7, Q10231, OSC) as queries. Eukaryotic and bacterial sequences are indicated by black and grey bars, respectively. The collapsed clade contains 37 leaves from plants and green algae. 100 bootstrap replicates were performed, values above 70 are shown on the corresponding branches. All sequences and the final tree are provided in SI Dataset S02 and SI Dataset S03, respectively. The tree was midrooted, visualized and made available in iTOL (https://itol.embl.de/tree/8384480491291613138765). (**A**) Distribution of HMMER E-values obtained with *Sch. japonicus* SHC (SCHJC_C003990) as query against a bacterial sequence database (grey bars) and a eukaryotic database (black bars). (**B**) Distribution of HMMER E-values obtained with *Sch. japonicus* OSC (SCHJC_A005630) as query against a bacterial sequence database (grey bars) and a eukaryotic database (black bars).

The systematic search for triterpene cyclase homologs and subsequent phylogenetic analysis showed that the putative *Sch. japonicus* SHC sequences (SCHJC_C003990 from CBS5679, and B6K412 from yFS275) are related to bacterial SHCs, with sequences from *Acetobacter* spp. (A0A0D6N754 from *A. indonesiensis* 5H-1, and A0A0D6NG57 from *A. orientalis* 21F-2) as closest relatives (Figure 4). To check if this conclusion was biased by the selection of sequences from species of interest, the putative SHC sequence SCHJC_C003990 from *Sch. japonicus* CBS5679 was used as query for a second HMMER search of either the eukaryotic or the bacterial databases described above. The resulting E-values distribution (Figure 4A; SI Dataset S05) showed a strong overrepresentation of low E-values among prokaryotic sequences. Sequences of two *Acetobacter* species, A0A0D6NG57 from *A. orientalis* 21F-2 and A0A0D6N754 from *A. indonesiensis* 5H-1, showed 67.9 % and 66.9 % sequence identity, respectively, and yielded zero E-values in this search. In contrast, E-value distributions obtained with the putative OSC sequence SCHJC_A005630 from *Sch. japonicus* CBS5679 as query showed an overrepresentation of low E-values among eukaryotic sequences (Figure 4B).

No SHC homologs were found in the predicted proteomes of Schizosaccharomycetes other than *Sch. japonicus*, nor in those of 371 Saccharomycotina yeast species included in the eukaryotic UniProt database. Acquisition of an STC-encoding DNA sequence from bacteria has been proposed as a key event in the evolution of strictly anaerobic fungi^46^. Since the putative *Sch. japonicus* SHC homologs did not show homology with STCs from those deep-branching fungi or from ciliates (Figure 4), acquisition of a bacterial SHC-encoding DNA sequence by an ancestor of *Sch. japonicus* could represent an independent eukaryotic adaptation to an anaerobic lifestyle.

### *Sch. japonicus* synthesizes hopanoids

Since SHCs can catalyze the conversion of squalene to various polycyclic triterpenoids^47,48^, we assessed whether the unidentified components in triterpenoid fractions of anaerobically grown *Sch. japonicus* (Figure 3B) were hopanoids. GC-MS analysis of biomass samples (Figure 3C-F; SI Appendix Table S4, SI Dataset S06) yielded 8 distinct analytes that were detected in *Sch. japonicus* (Figure 3B) but not in *S. cerevisiae* CEN.PK113-7D (Figure 3A). Squalene epoxide (compound **3**) was identified based on relative retention time and spectral matching with authentic standard material as previously described^49^, and hop-22(29)-ene (diploptene, compound **12**) was identified similarly (Figure 3D). For the remaining six compounds, structures were explored based on published mass spectra of hopanoids^50^ (Figure 3; SI Appendix Table S4, SI Dataset S06). A fragment ion with a mass-charge ratio (*m/z*) of 191, which is present in mass spectra of many hopanoids and frequently as the base peak^50^, was detected for compounds **4**, **13** and **14** (Figure 3C, E and F) and comparison with published data putatively identified them as hop-17(21)-ene, hop-21(22)-ene and hopan-22-ol (diplopterol), respectively^11,50^. Mass and retention-time shifts caused by silylation^51^ were investigated (Figure 3) and confirmed presence of the hydroxy group of diplopterol (Figure 3B; **14** and **15**) in the *Sch. japonicus* biomass, as well as those of the sterols (Figure 3A; **6** and **7**, **10** and **11**) in the *S. cerevisiae* samples. A small peak at the retention time of unsilylated diplopterol was tentatively attributed to steric hindrance by the tertiary-alcohol context of its hydroxy group. In the chromatograms representing silylated triterpenoids of *S. cerevisiae*, small additional peaks at retention times of 16.0 min and 16.6 min were attributed to ergosta-5,7,22-trien-3β-ol (also detected in the commercial ergosterol preparation used for supplementation of anaerobic growth media; SI Appendix, Figure S2) and fecosterol, an intermediate of ergosterol biosynthesis^52^, respectively (Figure 3A).

Compound **5** showed the characteristic base peak of *m/z* 191 and a molecular ion of *m/z* 428, pointing towards a hydroxylated structure analogous to diplopterol, but was not readily silylated. For substances **8** and **9**, the ion at *m/z* 410 suggests an unsubstituted triterpene which is unaffected by silylation, with retention times similar to those of other identified hopenes (Figure 3B). The corresponding base peaks of *m/z* 259 and 243 were previously reported for polycyclic triterpenoids with a different backbone configuration than those of substances **4**, **12** and **13**^53^. However, the precise structure of compounds **5**, **8** and **9** could not be identified with a high degree of certainty. All acquired mass spectra are shown in SI Dataset S06.

The newly identified compounds were quantified by GC-FID analysis in biomass samples of anaerobic *Sch. japonicus* cultures supplemented with different combinations of Tween 80 and ergosterol. The detection of ergosterol in biomass from anaerobic cultures grown in the presence of this compound indicated that *Sch. japonicus* is able to import sterols (Figure 5A). Except for the presence and absence of ergosterol, the triterpenoid composition of anaerobically grown biomass was not markedly affected by the supplementation of ergosterol and/or Tween 80. In addition, analyses were performed on aerobically grown *Sch. japonicus* to investigate whether hopanoid synthesis in this yeast is affected by oxygen availability. This experiment confirmed the ability of *Sch. japonicus* strain CBS5679 to synthesize ergosterol (Figure 5A). Aerobically grown biomass showed a 3.5-fold higher squalene content than biomass grown in anaerobic cultures without Tween 80 and ergosterol, while its hopanoid content was 4-fold lower (Figure 5A; SI Dataset S01). These observations suggest that oxygen availability may regulate triterpenoid synthesis in *Sch. japonicus*.

**Figure 5.**
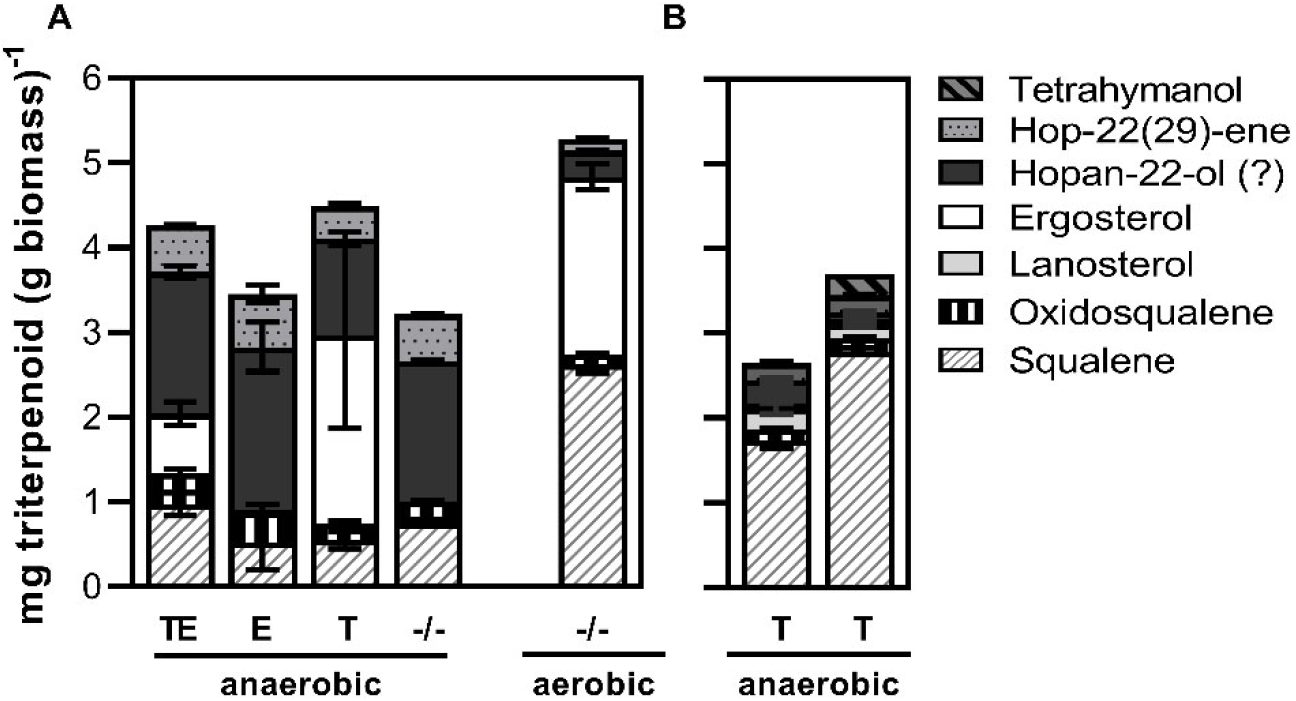
Quantification of triterpenoids in yeast biomass. (**A**) Triterpenoid content of cultures of *Sch. japonicus CBS5679* grown in SMPD with 20 g L^−1^ glucose. Under anaerobic conditions, cultures were supplemented with Tween 80 and ergosterol (TE), only ergosterol (E), only Tween 80 (T) or neither of these supplements (−/−). Data are shown for the first anaerobic culture following the anaerobic pre-culture. Aerobic cultures of *Sch. japonicus* were grown in SMPD without supplements (−/−). (**B**) Triterpenoid composition of anaerobic cultures of *S. cerevisiae* IMX2616 (*sga1Δ::Sjshc1*; left) and IMX2629 (*sga1Δ::Sjshc1 X-2::Maths*; right) grown in SMPD with 20 g L^−1^ glucose and Tween 80 (T) supplementation. Data are represented as average ± SEM of data from two independent duplicate cultures for each cultivation condition. Detailed information on data presented in this figure and additional anaerobic transfers are provided in SI Dataset S01.

### Expression of *Sch. japonicus* squalene-hopene cyclase supports sterol-independent anaerobic growth of *S. cerevisiae*

To investigate if the putative squalene-hopene-cyclase gene of *Sch. japonicus* CBS5679 (*Sjshc1*) was responsible for hopanoid synthesis, its coding sequence was codon optimized and expressed in the Cas9-expressing *S. cerevisiae* strain IMX2600. Growth and triterpenoid production of the resulting strain IMX2616 (*sga1Δ::Sjshc1*; SI Appendix, Table S6) was studied in anaerobic shake-flask cultures. After an anaerobic pre-culture for depletion of cellular reserves of sterols and/or hopanoids, neither the reference strain *S. cerevisiae* CEN.PK113-7D nor the strain carrying the *Sjshc1* expression cassette grew on SMPD without ergosterol and Tween 80 (Figure 6). On SMPD with only Tween 80, *S. cerevisiae* CEN.PK113-7D reached an optical density of 0.7 after 33 h (Figure 6A), at which point approximately 70 % of the initially present glucose remained unused (SI Appendix; Table S5). In contrast, *S. cerevisiae* IMX2616 (*sga1Δ::Sjshc1*) reached an optical density of 2.1 after the same time period (Figure 6B), at which point 98 % of the initially added glucose had been consumed (SI Appendix; Table S5), and showed sustained anaerobic growth upon transfer to a second flask containing the same medium (Figure 6B). Upon termination of the experiments after 58 h, the OD_600_ of the *S. cerevisiae* CEN.PK113-7D cultures had increased to 1.1. Similar very slow growth in sterol-free media was previously attributed to minute oxygen leakage^20^.

**Figure 6.**
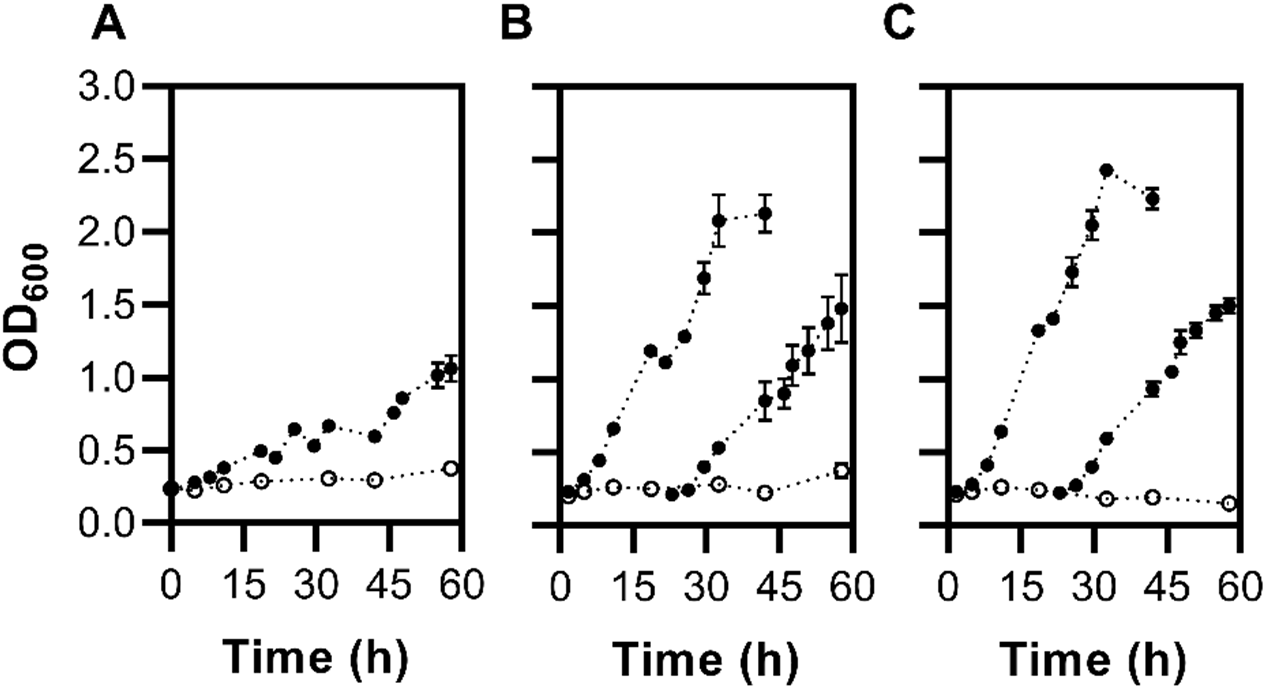
Anaerobic growth of *S. cerevisiae* strains in sterol-free media. *S. cerevisiae* cultures were inoculated from an anaerobic pre-culture on SMPD (50 g L^−1^ glucose) to fresh SMPD (20 g L^−1^ glucose), either supplemented with Tween 80 (closed circles), or lacking unsaturated fatty acids and sterols (open circles). (**A**) Reference strain CEN.PK113-7D. (**B**) *S. cerevisiae* strain IMX2616 (*sga1Δ::Sjshc1*). (**C**) *S. cerevisiae* strain IMX2629 (*sga1Δ::Sjshc1 X-2::Maths*). Cultures supplemented with Tween 80 represented in panel (**B**) and (**C**) were transferred to fresh medium of the same composition (closed circles) during exponential phase. Data are represented as average ± SEM of measurements on independent duplicate cultures for each yeast strain.

To investigate whether *S. cerevisiae* IMX2616 (*sga1Δ::Sjshc1*) produced the same hopanoid compounds as *Sch. japonicus* CBS5679, biomass was harvested from anaerobic shake-flask cultures grown on SMDP with Tween 80. Analysis of the triterpenoid fraction by GC-MS and GC-FID showed the same hopanoids that were detected and identified in *Sch. japonicus* strain CBS5679 (SI Appendix; Figure S3), albeit in smaller amounts (Figure 5B; SI Dataset S01). The only sterol identified in these samples was lanosterol. Synthesis of small quantities of this first tetracyclic intermediate of ergosterol biosynthesis by *S. cerevisiae* strains has been attributed to small oxygen leakages into anaerobic cultivation systems^20^.

In some prokaryotes, SHC is involved in a pathway for tetrahymanol production, in which a tetrahymanol synthase converts hopene into tetrahymanol^11^. To investigate whether such a two-step pathway for tetrahymanol synthesis can be engineered in *S. cerevisiae*, a codon-optimized expression cassette for the *Methylomicrobium alcaliphilum*^11^ gene encoding THS (locus tag MEALZ_1626; referred to as *Maths*) was integrated at the X-2 locus^54^ in strain IMX2616 (*sga1Δ::Sjshc1*), yielding strain IMX2629 (*sga1Δ::Sjshc1 X-2::Maths*; SI Appendix, Table S6). Anaerobic growth and sugar consumption rates of these two *S. cerevisiae* strains were similar (Figures 6B and 6C; SI Appendix, Table S5). Tetrahymanol was detected in anaerobically grown biomass of strain IMX2629, but not of strain IMX2616 (Figure 5C; SI Appendix, Figure S4). Together, these results confirm that *Sjshc1* encodes a bona fide SHC, at least one of whose hopanoid products can act as sterol surrogate in anaerobic yeast cultures.

## Discussion

Research on obligately anaerobic fungi belonging to the phylum Neocallimastigomycota has dispelled the notion that sterols are indispensable components of all eukaryotic membranes. Growth studies linked the ability of these fungi to maintain proper membrane function in the absence of oxygen or exogenous sterols to their ability to synthesize the triterpenoid sterol surrogate tetrahymanol^41,46^. A recent study on expression of squalene-tetrahymanol cyclase in the model eukaryote *S. cerevisiae*^19^ confirmed that tetrahymanol synthesis is not only required but also sufficient for sterol-independent growth. Inspired by a half-century old, intriguing publication by Bulder^21^ on the yeast *Sch. japonicus*, the present study uncovered a different eukaryotic adaptation to circumvent oxygen requirements for sterol biosynthesis.

GC-MS analysis identified several hopanoids in anaerobically grown *Sch. japonicus* CBS5679 (Figure 3; SI Appendix, Table S4) that were subsequently also detected in an *S. cerevisiae* strain expressing a codon-optimized *Sch. japonicus* ORF with sequence homology to known prokaryotic squalene hopene cyclase (SHC) genes (Figure 6; SI Appendix, Figure S4). These results indicate that product diversity originated from the *Sch. japonicus Sjshc1*-encoded SHC, rather than from additional enzyme-catalyzed modifications. Formation of multiple products is consistent with reports on triterpenoid extracts of bacterial hopanoid producers and product spectra of purified bacterial SHCs^55,56^. For example, analysis of triterpenoids in *Zymomonas mobilis* biomass revealed a number of minor hopene variants, in addition to diploptene and diplopterol, whose synthesis was attributed to deviation from the regular cyclization process^57^.

Functional SHC enzymes and hopanoid synthesis have been found in ferns^58,59^ and putative SHC proteins have been identified in several filamentous fungi^60,61^ (Figure 4). Although *Sch. japonicus* is therefore not unique among eukaryotes in containing an SHC, hopanoid synthesis has not previously been found in yeasts or associated with sterol-independent anaerobic growth of eukaryotes. No putative SHC genes were found in other *Schizosaccharomyces* species, nor in more distantly related yeasts. Confinement to a single yeast species and a strong similarity with putative SHC sequences from *Acetobacter* species identifies horizontal gene transfer (HGT) from acetic acid bacteria as a highly plausible evolutionary origin of *Sjshc1*^60^. Similar HGT events have been implicated in acquisition of squalene-tetrahymanol cyclase during transitions of eukaryotic lineages from aerobic to anaerobic lifestyles^46,60^. The independent acquisition of different squalene-cyclase genes by phylogenetically distant eukaryotes represents a remarkable case of convergent evolution towards an anaerobic eukaryotic lifestyle.

In the absence of efficient procedures for genetic modification of *Sch. japonicus* CBS5679^62^, the role of one or more hopanoids as sterol surrogates was confirmed by heterologous expression of *Sjshc1* in *S. cerevisiae*, which enabled anaerobic growth under ergosterol-depleted conditions (Figure 6). This positive impact on growth of a yeast in an anaerobic, sterol-free environment illustrates how the mere acquisition of an SHC gene by HGT may have benefited an ancestor of *Sch. japonicus*. However, ergosterol-independent anaerobic growth of the *Sjshc1*-expressing *S. cerevisiae* strain, as well as that of a strain that co-expressed *Sjshc1* with a bacterial tetrahymanol synthase from *M. alkaliphilum*^11^ was much slower than that of *Sch. japonicus* and of ergosterol-supplemented *S. cerevisiae* cultures^37^ (Figure 6). A reduced specific growth rate in sterol-free media was previously also observed for *S. cerevisiae* and *Kluyveromyces marxianus* strains expressing a squalene-tetrahymanol cyclase gene from the ciliate *T. thermophila*^20,63^.

Our results confirm the report by Bulder^21^ that *Sch. japonicus* does not require supplementation of unsaturated fatty acids (UFAs) for anaerobic growth. However, we did not find evidence for his hypothesis that this yeast is capable of oxygen-independent UFA synthesis^39^. Instead, in comparison with UFA-supplemented cultures, cultivation in UFA-free medium led to a reduction of the average chain length of saturated fatty acids (SFA; Figure 2). A similar shift in SFA composition has been observed during slow UFA-independent anaerobic growth of *S. cerevisiae* CEN.PK113-7D^37^ and in membranes of anaerobically grown cultures of the dimorphic fungus *Mucor rouxii*^64^.

Our results indicate that its membrane composition and/or membrane architecture enable *Sch. japonicus* to maintain a much faster growth rate than *S. cerevisiae* upon sterol replacement by hopanoids, as well as in the absence of UFAs. A recent study on membrane properties of *Sch. japonicus* and the more closely related yeast *Sch. pombe* showed a higher membrane stiffness and lipid packing in *Sch. japonicus* as well as generally shorter and more saturated fatty acid residues than found in *Sch. pombe*^33^. Further research should resolve the impact of hopanoids on membrane properties of *Sch. japonicus*. In bacteria, hopanoids have been implicated in tolerance to external stresses such as non-optimal temperature and pH, and the presence of antimicrobials^65–67^. Elucidating how sterol surrogates interact with other membrane components, including proteins, and thereby influence membrane functionality can contribute to a deeper insight in microbial adaptation to anaerobic environments and to physicochemical stress factors. In addition, such studies will contribute to the design of membrane engineering strategies aimed at the construction of robust industrial strains of *S. cerevisiae* and other yeasts for application in anaerobic fermentation processes.

## Materials and methods

### Strains, media and maintenance

*Schizosaccharomyces japonicus* CBS5679 was obtained from the Westerdijk Institute (Utrecht, The Netherlands). *Saccharomyces cerevisiae* strains used and constructed in this study belonged to the CEN.PK lineage^68,69^ (SI Appendix, Table S5). Yeast strains were propagated in YPD and stored at −80°C after addition of 30 % sterile glycerol^20^. To avoid sexual co-flocculation^70^ of *Sch. japonicus* CBS5679, carbon source depletion in pre-cultures was prevented and buffered synthetic media were used in all growth studies. Yeasts were grown on synthetic medium with ammonium as nitrogen source^71^ with an increased concentration of KH_2_PO_4_ (14.4 g L^−1^, SMP^72^). Unless otherwise indicated, glucose was added from a concentrated stock solution, separately autoclaved at 110°C, to a concentration of 20 g L^−1^ (SMPD). Where indicated, SMPD was supplemented with Tween 80 (polyethylene glycol sorbitan monooleate; Merck, Darmstadt, Germany) and/or ergosterol (≥95 %; Sigma-Aldrich, St. Louis, MO) at concentrations of 420 mg L^−1^ and 10 mg L^−1^, respectively, from a sterile 800-fold concentrated stock solution^37^. Bacto Agar (BD Biosciences, 20 g L^−1^) was added to prepare solid media. The counter-selectable *amdS*-marker was used as described previously^73^. Strains with geneticin, hygromycin or nourseothricin resistance were selected by supplementing YPD with 200 mg L^−1^ geneticin (G418), 100 mg L^−1^ hygromycin B (hygB) or 100 mg L^−1^ nourseothricin (ClonNAT), respectively.

### Molecular biology techniques

Open-reading frames of a putative squalene-hopene-cyclase gene (SHC; SCHJC_C003990) from *Sch. japonicus* CBS5679 and of a *Methylomicrobium alkaliphilum* tetrahymanol synthase gene (Genbank Accession number CCE23313) were codon-optimized for use in *S. cerevisiae* with the GeneOptimizer algorithm (GeneArt, Regensburg, Germany)^74^. DNA fragments for plasmid construction were amplified with Phusion High-Fidelity DNA Polymerase (Thermo Scientific, Waltham, MA) as specified by the manufacturer, using PAGE-purified oligonucleotide primers (Sigma-Aldrich). Diagnostic PCR was performed with DreamTaq PCR Master Mix (Thermo Scientific), according to the manufacturer’s protocol using desalted oligonucleotides (Sigma-Aldrich). PCR-amplified linear integration cassettes were purified from 1 % (w/v) agarose gels (TopVision Agarose, Thermo Fisher) with TAE buffer (Thermo Fisher) using a Zymoclean Gel DNA Recovery Kit (Zymo Research, Irvine, CA). Assembly of DNA fragments was performed with Gibson Assembly Master Mix (New England Biolabs, Ipswich, MA). *E. coli* XL^−1^ Blue cells (Agilent Technologies, Santa Clara, CA) were transformed with assembled plasmids according to the provider’s instructions and stored at −80 °C. *S. cerevisiae* was transformed with the lithium-acetate method^75^ and marker-less CRISPR/Cas9-based genome editing of *S. cerevisiae* was performed as described previously^76^. Transformants were selected on YPD-CloNAT, YPD-G418 or YPD-hygB agar for IMX2600, IMX2616 and IMX2629 respectively with subsequent isolation of single-cell lines by three consecutive re-streaks. Plasmids and oligonucleotides used are listed in SI Appendix, Tables S7 and S8, respectively.

### Plasmid and strain construction

All *S. cerevisiae* strains that were used and constructed in this study are listed in SI Appendix, Table S6. A *S. cerevisiae* strain suitable for marker-less genome editing^76^ was constructed by integrating of expression cassettes for *Spcas9* and the *natNT2* marker gene in the *CAN1* locus of *S. cerevisiae* CEN.PK113-7D. Transformation of CEN.PK113-7D with 2000 ng and 900 ng of the *Spcas9* and *natNT2* cassettes respectively, yielded strain IMX2600. These integration cassettes were amplified from p414-TEF1p-cas9-CYC1t^77^ and pUG-natNT2^78^, respectively, using primer pairs 2873/4653 and 3093/5542. Expression cassettes for the putative SHC-gene of *Sch. japonicus* (*Sjshc1*) and the THS-gene from *M. alcaliphilum* (locus tag MEALZ_1626; referred to as *Maths*) used the *S. cerevisiae TEF1* promoter and *CYC1* terminator, or *TDH3* promoter and *ADH1* terminator sequences, respectively. Codon-optimized coding sequences were first amplified from plasmids pUD1151 and pUD1150 using primer pairs 15100/15101 and 17519/17520, respectively. The p426TEF and pUD63 backbones were linearized using primer pairs 5921/10547 and 10546/3903, respectively. Gibson assembly of the linearized p426TEF-backbone and *Sjshc1*-insert resulted in plasmid pUD1059, and assembly of the pUD63-backbone and *Maths*-insert resulted in plasmid pUDE1060. The expression cassettes for the *Sjshc1*-gene and *Maths*-gene were then amplified from these plasmids using primer pairs 15002/15003 and 9034/17521, respectively. The *Sjshc1-*construct was integrated in the *SGA1* locus of *S. cerevisiae* IMX2600 by co-transformation with 500 ng of the linear fragment and 500 ng of the *SGA1*-targeting plasmid pUDR119, resulting in *S. cerevisiae* IMX2616. Correct integration was verified by diagnostic PCR using primers 7298/7479 (both binding outside the *SGA1* locus) and 7298/15103 and 7479/15102 (binding outside of the integration locus and inside of the heterologous SHC-gene). To target the X-2 locus, pUDR538 was constructed by Gibson assembly of the backbone of pROS12, linearized with primer 6005, and a 2μm cassette amplified from pROS12 with primer 10866. The *Maths*-gene was subsequently integrated in the X-2 locus^54^ of *S. cerevisiae* strain IMX2616 by co-transformation with 500 ng of the linear cassette and 500 ng of the X-2 targeting plasmid pUDR538, resulting in strain IMX2629. Correct integration was verified by diagnostic PCR using primers 7376/7377 (both binding outside the X-2 locus) and 7376/17523 and 7377/17522 (binding outside of the integration locus and inside of the heterologous THS gene).

### Aerobic shake-flask cultivation

Aerobic cultivation on SMPD was performed in 500-mL shake flasks with a working volume of 100 mL. A pre-culture was inoculated from a frozen stock culture and, after overnight incubation, transferred to a second pre-culture. Upon reaching mid-exponential phase, cells were transferred to fresh SMPD and optical density was monitored at 660 nm. All experiments were performed in duplicate. Light-induced flocculation of *Sch. japonicus*^79^ was prevented by wrapping flasks in aluminium foil.

### Anaerobic shake-flask cultivation

Anaerobic chamber experiments were performed as previously described^37^, using 100-mL shake flasks containing 80 mL of medium. The anaerobic chamber was placed in a mobile darkroom, which was only illuminated with red LEDs^80^ during sampling. An aerobic pre-culture on SMPD was used to inoculate an anaerobic pre-culture on SMPD with an increased glucose concentration of 50 g L^−1^, which was grown until the end of the exponential phase. Samples from these pre-cultures were then used to inoculate experiments on regular SMPD supplemented with either Tween 80 and ergosterol, only Tween 80 or ergosterol, or neither. All growth experiments were performed by monitoring optical density at 600 nm in independent duplicate cultures.

### Analytical methods

High-performance liquid chromatography was used to analyze extracellular metabolite concentrations as described previously^81^. Extraction of fatty acids and triterpenoids and quantitative analysis by gas-chromatography with flame ionization detection (GC-FID) was performed as previously described^20,37^. The GC-FID system was calibrated for hop-22(29)-ene (Sigma Aldrich, 0.1 mg mL^−1^ in isooctane) using a 6-point calibration, and this calibration curve was additionally used for quantification of other detected hopanoid compounds. For anaerobic cultures, optical density at 600 nm was measured with an Ultrospec 10 cell density meter (Biochrom, Harvard Biosience, Holliston, MA) placed in the anaerobic chamber. In aerobic cultures, optical density at 660 nm was measured on a Jenway 7200 spectrophotometer (Bibby Scientific, Staffordshire, UK).

For gas chromatography-mass spectrometry (GC-MS) analysis, a Varian 3800 gas chromatograph with a CombiPal autosampler (CTC Analytics, Zwingen, Switzerland) was coupled with a Saturn 2200 ion trap MS with a Varian 1177 injector set in 1:3 split mode (Varian, Darmstadt, Germany) controlled by Varian Workstation 6.9 SP 1 software. An Agilent VF 5ms (Agilent Technologies) capillary column of 30 m length with a 10 m EZ Guard column (0.25 mm internal diameter and 0.25 μm film thickness) was used with an inlet temperature of 250 °C, and an injection volume of 1 μL (splitless time 1.0 min). Helium (99.999%; Air Liquide, Düsseldorf, Germany) was used as carrier gas at a constant flow rate of 1.4 mL min-1. The GC oven started at 55 °C (1.0 min hold), ramped up to 260 °C at 50 °C min-1, followed by a gradient of 4 °C min-1 up to 320 °C (hold time 3.9 min). The total run time was 24.0 min. The transfer line temperature was set at 270 °C and the ion-trap temperature was 200 °C. The ion trap was operated in two segments. The mass spectrometer (MS) was switched off (solvent delay) for the first 10 min, and from 10 to 24 min the MS scanned at a mass range from 50 to 600 m/z (EI, 70 eV). Data analyses were carried out with the Agilent MassHunter Workstation Software package B.08.00 (Agilent Technologies). Triterpenoid compounds were identified by comparison with commercial references^20,51^ or literature data^11,50^. Autosampler vials and caps and the silylation reagents N-trimethylsilylimidazole (TSIM) and N-methyl-N-trimethylsilyltrifluoroacetamide (MSTFA) were purchased from Macherey-Nagel (Düren, Germany). The primary secondary amine (PSA) for dispersive solid phase extraction (dSPE) was acquired from Agilent Technologies. Ergosterol (≥ 95.0%), 5α-cholestane (≥ 97.0%) and hop-22(29)-ene (0.1 mg/mL in isooctane, analytical standard) were obtained from Sigma-Aldrich. Other reagents and solvents were purchased in HPLC-grade or pro-analysis quality from Sigma-Aldrich and all consumables were from VWR (Ismaning, Germany). 8 mg of freeze-dried yeast biomass was mixed with 2 M NaOH to obtain a final concentration of 4 mg biomass mL^−1^. Subsequent procedures were performed as described previously^51^, with slight modifications. After saponification, the suspension was divided in two microcentrifuge tubes (2 × 500 μL). 650 μL of distilled methyl-tert-butylether (MtBE) and 100 μL of internal standard solution (5α-cholestane in MtBE, 10 μg mL^−1^) were added to each sample. The mixtures were shaken well for 1 min and centrifuged at 9000 × *g* for 5 min. The organic upper layer was transferred into a new 2.0 mL plastic microcentrifuge tube containing 40 ± 2 mg of a mixture (7:1) of anhydrous sodium sulfate and primary secondary amine (PSA). The mixture was extracted a second time in the same manner with another 750 μL of MtBE. The combined organic extracts were vigorously shaken for 1 min, followed by a centrifugation step (5 min, 9000 × *g*). 1 mL of the purified upper layer was then transferred into a brown glass vial and concentrated to dryness under a stream of nitrogen. One residue was dissolved in 700 μL of MtBE and 50 μL of silylation reagent mixture MSTFA/TSIM (9:1) was added. The other residue was dissolved in 750 μL of MtBE before analysis.

### Genome sequencing and assembly

The genome of *Sch. japonicus* CBS5679 was sequenced using short-read and long-read sequencing technologies. Genomic DNA was isolated with a Qiagen genomic DNA 100/G kit (Qiagen, Hilden, Germany) with modifications to the ‘Part I sample preparation and lysis protocol for yeast’ of the manufacturer’s instructions. Yeast cells were harvested by centrifugation (10 min at 3700 × *g*) from 100 mL overnight cultures on YPD. Zymolyase incubation was replaced by freezing yeast pellets in liquid nitrogen, manual grinding and resuspension in buffer G2 with RNAse A. Incubation with Proteinase K was extended to 3 h. Further steps were as described in the manufacturer’s protocol. DNA quantity was measured on a Qubit Fluorometer (Invitrogen, Carlsbad, CA) using a QuBit BR dsDNA Assay kit (Invitrogen). For short read sequencing, 150 bp paired-end libraries were prepared with a Nextera DNA Flex Library Prep kit (Illumina, San Diego, CA) according to the manufacturer’s instructions and whole-genome sequencing was performed on an in-house MiSeq platform (Illumina). For long-read sequencing, genomic libraries were prepared using the 1D Genomic DNA by ligation (SQK-LSK109, Oxford Nanopore Technologies, Oxford, UK) according to the manufacturer’s instructions, with the exception of the ‘End Repair/dA-tailing module’ step during which the ethanol concentration was increased to 80 % (v/v). Quality of flow cells (R9 chemistry flow cell FLO-MIN106, Oxford Nanopore Technologies) was tested with the MinKNOW platform QC (Oxford Nanopore Technologies). Flow cells were prepared by removing 20 μL buffer and subsequently primed with priming buffer. The SQK-LSK109 DNA library was loaded dropwise into the flow cell and sequenced for 48 h. Base-calling was performed using Guppy v2.1.3 (Oxford Nanopore Technologies) using dna_r9.4.1_450bps_flipflop.cfg. Genome assembly was performed using Flye v2.7.1-b167359^82^. Flye contigs were polished using Pilon v1.18^83^. The polished assembly was annotated with Funannotate v1.7.1^84^ using RNAseq data from bioprojects PRJNA53947 and PRJEB30918 as evidence of transcription, adding functional information with Interproscan v5.25-64.0^85^.

### Sequence homology search and phylogenetic analyses

Eukaryotic and bacterial amino acid sequence databases were built from UniProtKB reference proteomes (Release 2019_02) using the taxonomic divisions (taxids) shared with databases from the National Center for Biotechnology Information (NCBI). Only amino acid sequences from reference or representative organisms having genome assemblies at chromosome or scaffold level according to the NCBI genomes database (Release 2019_03) were included. Table S3 (SI Appendix) indicates proteomes from species of interest, which were included in the corresponding databases obtained from the UniProt TrEMBL division. The *Sch. japonicus* CBS5679 proteome was obtained in this study and consequently not available from UniProt at the time of analysis. Amino acid sequences of an oxidosqualene cyclase (OSC) from *Sch. pombe*^42^ (SpErg7; UniProt accession Q10231), a squalene-hopene cyclase (SHC) from *Acidocaldarius alicyclobacillus*^43^ (AaShc; P33247), and a squalene-tetrahymanol cyclase (STC) from *Tetrahymena thermophila*^5,20^ (TtThc1; Q24FB1) were used as queries for a HMMER3^44^ homology search. HMMER hits with an E-value below 1×10^−5^ and a total alignment length (query coverage) exceeding 75 % of the query sequence were considered significant. The taxids of 28 eukaryotic and 20 bacterial species of interest (SI Appendix, Table S3) were used to select sequences from all significant bacterial and eukaryotic HMMER hits obtained using the three cyclase sequences as queries. This procedure yielded 128 sequences (45 bacterial, 83 eukaryotic) including the 10 HMMER hits with the lowest E-values obtained by using either SjErg7 or SjShc1 as queries against the bacterial and eukaryotic databases (SI Dataset S02). These 128 sequences were subjected to multiple sequence alignment using MAFFT v7.402^86^ in “einsi” mode. Alignments were trimmed using trimAl v1.2^87^ in “gappyout” mode, and used to build a phylogenetic tree with RAxML-NG v0.8.1^88^ using 10 random and 10 parsimony starting trees, 100 Felsestein Bootstrap replicates, and PROTGTR+FO model. The final, midrooted tree provided in SI Dataset S03 was visualized using iTOL^89^.

## Data availability

Whole-genome sequencing data for *Sch. japonicus* CBS5679 have been deposited under the BioProject accession PRJNA698797 in NCBI.

## Acknowledgements

This work was funded by an Advanced Grant of the European Research Council to JTP (grant 694633). We gratefully acknowledge Jasmijn Hassing for constructing strain IMX2600, and Nicolo Baldi for the construction of plasmid pUDR538. We thank our colleagues in the Industrial Microbiology group of TU Delft for stimulating discussions.

