## Supplementary material for "A squalene-hopene cyclase in *Schizosaccharomyces japonicus* represents a eukaryotic adaptation to sterol-independent anaerobic growth": SI Appendix

[1] Shared first authors

**This PDF file contains:**

Figures S1-S5

Tables S1-S8

**Other supplementary materials for this manuscript include the following:**

Datasets S01-S06

**Supplementary Figures**

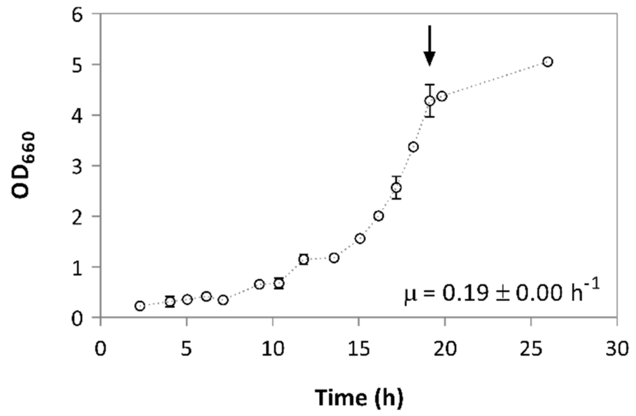


**Fig. S1. Aerobic growth of *Sch. japonicus* CBS5679.** An exponentially growing aerobic pre-culture on SMPD (20 g $L^{-1}$ glucose) was used to inoculate a 100 mL-culture on the same medium in 500 mL shake-flasks. Flasks were wrapped in aluminum foil to avoid light-induced sexual flocculation[1,2]. Optical density at 660 nm was measured over time for two biological duplicates, and data are presented as the mean and average deviation. Specific growth rate (μ) was calculated from 11 points in the exponential phase. The arrow indicates the time point at which biomass was harvested for analysis by gas chromatography-mass spectrometry (GC-MS).

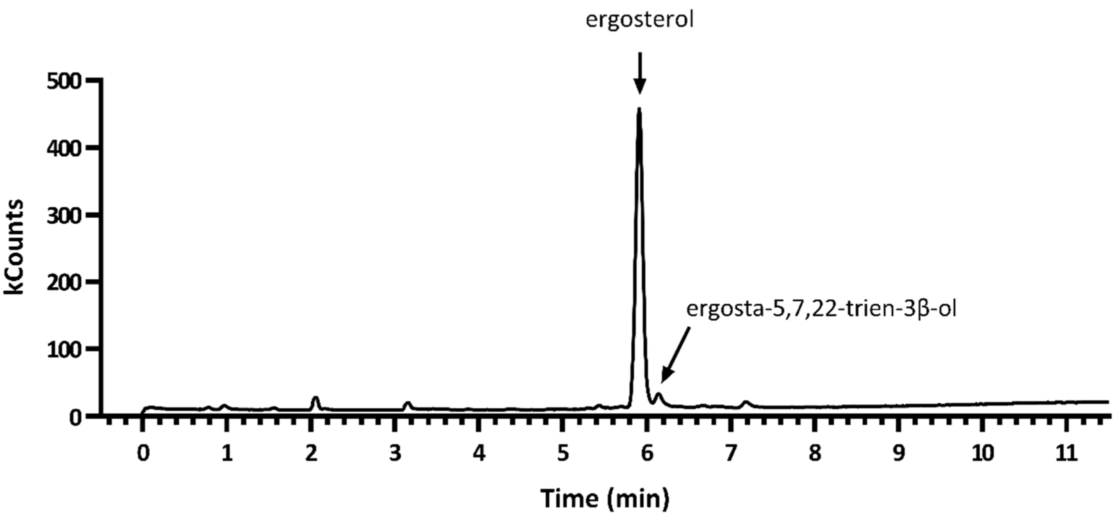


**Fig. S2. GC-MS analysis of commercial ergosterol preparation.** A sample of the commercial ergosterol preparation (≥95% pure, Sigma-Aldrich) used for medium supplementation was subjected to the same work-up procedure used to analyze triterpenoid fractions of yeast biomass samples. The figure shows a chromatogram of the subsequent GC-MS analysis. Ergosterol and ergosta-5,7,22-trien-3β-ol were the only detected substances.

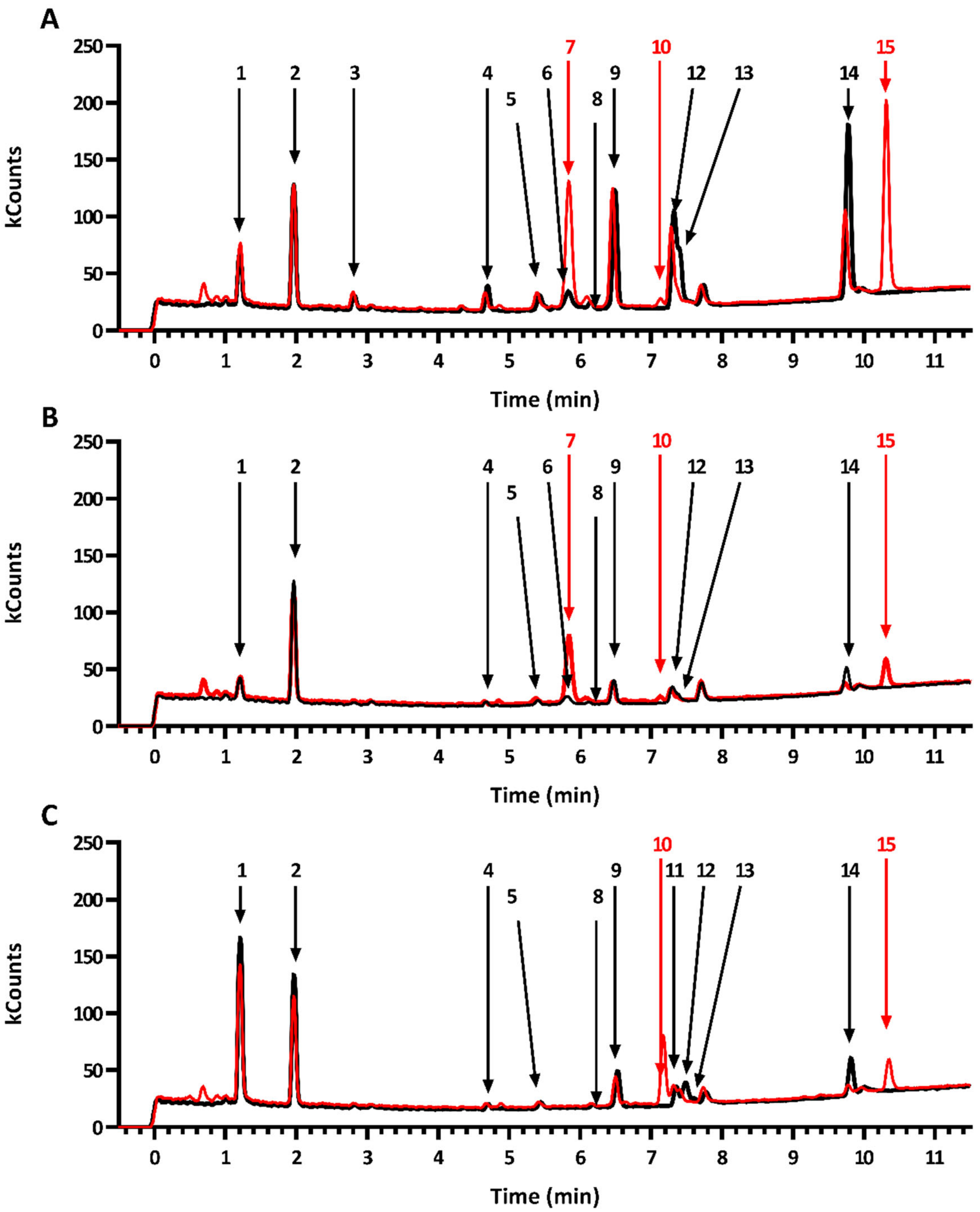


**Figure S3. Additional GC-MS chromatograms of the triterpenoid fraction of anaerobic yeast biomass.** Triterpenoids were extracted for GC-MS analysis from biomass harvested at the end of the exponential phase and either immediately injected (black lines) or first silylated (red lines). (**A**) Triterpenoid fraction of anaerobic biomass from *Sch. japonicus* strain CBS5679, grown on media supplemented with Tween 80 and ergosterol. (**B-C**) Triterpenoid fraction of anaerobic biomass from *S. cerevisiae* strain IMX2616 (*sga1Δ::SjSHC*) grown on media supplemented with

Tween 80 and ergosterol (**B**) or supplemented with only Tween 80 (**C**). Numbers indicate the following: **1**, squalene; **2**, 5α-cholestane (internal standard); **3**, squalene epoxide; **4**, hop-17(22)-ene; **5**, unidentified component; **6**, ergosterol; **7**, ergosterol-TMS-ether; **8**, unidentified component; **9**, unidentified component; **10**, lanosterol-TMS-ether; **11**, lanosterol; **12**, hop-22(29)-ene (diploptene); **13**, hop-21(22)-ene; **14**, hopan-22-ol (diplopterol); **15**, hopan-22-ol-TMS-ether.

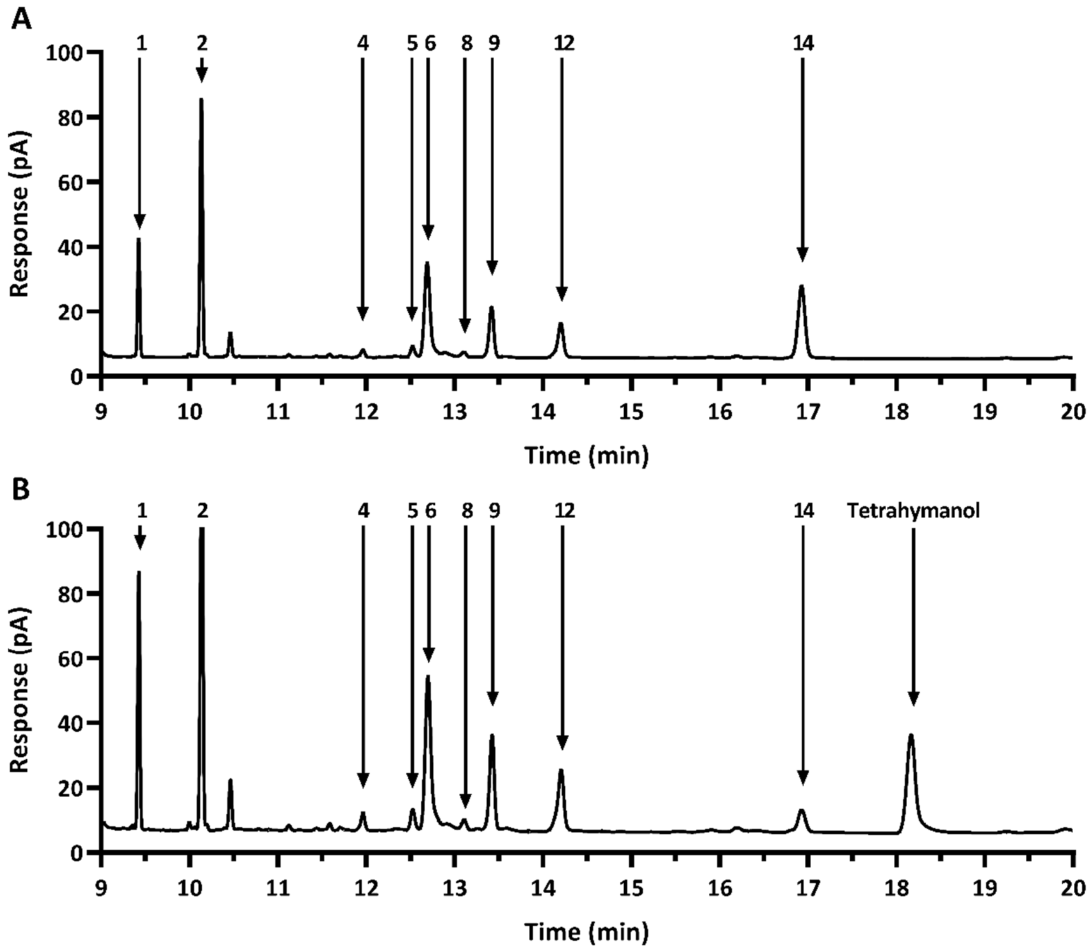


**Figure S4. GC-FID chromatograms of the triterpenoid fraction of anaerobic biomass of engineered *S. cerevisiae* strains.** Biomass was grown on media supplemented with Tween 80 and ergosterol, and harvested at the end of the exponential phase for triterpenoid extraction and subsequent GC-FID analysis. (**A**) Triterpenoid fraction of anaerobic biomass of *S. cerevisiae* strain IMX2616 (*sga1Δ::SjSHC*). (**B**) Triterpenoid fraction of anaerobic biomass of *S. cerevisiae* strain IMX2629 (*sga1Δ::SjSHC X-2::Maths*). Numbers indicate the following compounds: **1**, squalene; **2**, 5α-cholestane (internal standard); **3**, squalene epoxide; **4**, hop-17(22)-ene; **5**, unidentified component; **6**, ergosterol; **8**, unidentified component; **9**, unidentified component, possibly a tricyclic intermediate; **12**, hop-22(29)-ene (diploptene); **14**, hopan-22-ol (diplopterol).

**Supplementary tables accompanying experimental data**

**Table S1. Residual glucose concentrations upon termination of anaerobic shake-flask experiments with *Sch. japonicus*.** HPLC measurements of the residual glucose were made upon termination of each individual anaerobic shake-flask experiment. All experiments were performed on phosphate-buffered synthetic medium (SMPD). The anaerobic pre-culture contained 50 g $L^{-1}$ glucose (~278 mM), and all other experiments initially contained 20 g $L^{-1}$ glucose (~111 mM). Cultures were supplemented with Tween 80 and ergosterol (TE), only with Tween 80 (T), only with ergosterol (E), or with neither of those anaerobic growth factors (--). Data represent average and mean deviation of two independent cultures, and correspond to data presented in Figure 1 in the main text.

| Organism | Experiment | Glucose concentration (mM) |
|---|---|---|
| *S. cerevisiae* CEN.PK113-7D | Anaerobic pre-culture | 151.77 ± 22.38 |
| | SMPD +TE | 0.38 ± 0.19 |
| *Sch. japonicus* CBS5679 | Anaerobic pre-culture | 31.36 ± 11.37 |
| | TE | 0.00 ± 0.00 |
| | T, first transfer | 0.00 ± 0.00 |
| | T, second transfer | 0.00 ± 0.00 |
| | T, third transfer | 0.00 ± 0.00 |
| | E, first transfer | 0.10 ± 0.03 |
| | E, second transfer | 0.00 ± 0.00 |
| | E, third transfer | 0.14 ± 0.04 |
| | --, first transfer | 0.21 ± 0.08 |
| | --, second transfer | 0.00 ± 0.00 |
| | --, third transfer | 0.06 ± 0.00 |

**Table S2: Specific growth rates and final $OD_{600}$ for anaerobic shake-flask experiments with *Sch. japonicus* CBS5679.** Cultures were grown anaerobically on synthetic medium with an increased glucose concentration (50 g $L^{-1}$ glucose) in the absence of sterols or UFAs until the end of the exponential phase. Cells were transferred to SMPD with 20 g $L^{-1}$ glucose, supplemented with Tween 80 and ergosterol (TE), Tween 80 only (T), ergosterol only (E) or none of these anaerobic growth factors (--). Estimations of specific growth rate (μ) were based on four optical density measurements in the exponential phase. The final $OD_{600}$ indicates optical density at 600 nm of the cultures immediately before harvesting biomass for further analyses. Data represents the average and mean deviation of two independent cultures, and correspond to data presented in Figure 1 in the main text.

| **Medium** | | **Specific growth rate ($h^{-1}$)** | **Final $OD_{600}$** |
|---|---|---|---|
| SMPD, 50 g $L^{-1}$ glucose | | 0.29 ± 0.01 | 6.8 ± 0.4 |
| SMPD, 20 g $L^{-1}$ glucose, TE | | 0.27 ± 0.00 | 5.5 ± 0.1 |
| SMPD, 20 g $L^{-1}$ glucose, T | - (first transfer) | 0.28 ± 0.01 | 5.1 ± 0.3 |
| | - (second transfer) | 0.28 ± 0.01 | 5.6 ± 0.0 |
| | - (third transfer) | 0.30 ± 0.00 | 5.0 ± 0.4 |
| SMPD, 20 g $L^{-1}$ glucose, E | - (first transfer) | 0.28 ± 0.01 | 4.7 ± 0.8 |
| | - (second transfer) | 0.22 ± 0.00 | 4.2 ± 0.2 |
| | - (third transfer) | 0.25 ± 0.00 | 4.0 ± 0.3 |
| SMPD, 20 g $L^{-1}$ glucose, -- | - (first transfer) | 0.26 ± 0.00 | 4.3 ± 0.3 |
| | - (second transfer) | 0.24 ± 0.00 | 4.8 ± 0.1 |
| | - (third transfer) | 0.25 ± 0.00 | 4.3 ± 0.1 |

**Table S3. Species of interest used to retrieve cyclase homologs used in Figure 4.** Species of interest were selected based on (1) inclusion in Takishita *et al.*[3], (2) occurrence in deep-branching fungi[4], (3) occurrence in Schizosaccharomycetes[5]. Unless otherwise indicated, proteomes were obtained from UniProt reference proteomes (refprot). Bacterial or eukaryotic sequences are indicated as 'bact' and 'euk', respectively. NA, not applicable.

| Category | Species name | Taxid | Source | Origin |
|---|---|---|---|---|
| 1 | *Bacillus subtilis* | 1423 | Refprot | bact |
| 1 | *Blastopirellula marina* | 124 | Refprot | bact |
| 1 | *Bradyrhizobium japonicum* | 375 | Refprot | bact |
| 1 | *Frankia sp.* CcI3 *(Frankia casuarinae)* | 106370 | Refprot | bact |
| 1 | *Frankia sp.* EAN1pec | 298653 | TrEMBL | bact |
| 1 | *Geobacillus thermodenitrificans* | 33940 | TrEMBL | bact |
| 1 | *Geobacter sulfurreducens* | 35554 | Refprot | bact |
| 1 | *Gluconobacter oxydans* | 442 | Refprot | bact |
| 1 | *Methylococcus capsulatus* | 414 | Refprot | bact |
| 1 | *Nitrobacter hamburgensis* | 912 | Refprot | bact |
| 1 | *Nitrosococcus oceani* | 1229 | Refprot | bact |
| 1 | *Nitrosomonas europaea* | 915 | Refprot | bact |
| 1 | *Pelobacter propionicus* | 29543 | Refprot | bact |
| 1 | *Plesiocystis pacifica* | 191768 | Refprot | bact |
| 1 | *Rhodopseudomonas palustris* BisA53 | 316055 | TrEMBL | bact |
| 1 | *Rhodospirillum rubrum* | 1085 | Refprot | bact |
| 1 | *Saccharopolyspora erythraea* | 1836 | Refprot | bact |
| 1 | *Streptomyces coelicolor* | 1902 | Refprot | bact |
| 1 | *Syntrophobacter fumaroxidans* | 119484 | Refprot | bact |
| 1 | *Thermosynechococcus elongatus* | 146786 | Refprot | bact |
| 2 | *Anaeromyces robustus* | 1754192 | TrEMBL | euk |
| 1 | *Arabidopsis thaliana* | 3702 | Refprot | euk |
| 1 | *Aspergillus fumigatus* | 746128 | TrEMBL | euk |
| 1 | *Aureococcus anophagefferens* | 44056 | TrEMBL | euk |
| 1 | *Chlamydomonas reinhardtii* | 3055 | Refprot | euk |
| 1 | *Cryptococcus neoformans* | 5207 | Refprot | euk |
| 1 | *Danio rerio* | 7955 | Refprot | euk |
| 1 | *Dictyostelium discoideum* | 44689 | TrEMBL | euk |
| 1 | *Homo sapiens* | 9606 | Refprot | euk |
| 1 | *Mus musculus* | 10090 | Refprot | euk |
| 1 | *Naegleria gruberi* | 5762 | Refprot | euk |
| 2 | *Neocallimastix californiae* | 1754190 | TrEMBL | euk |
| 1 | *Neosartorya fischeri (Aspergillus fischeri)* | 36630 | Refprot | euk |

| | | | | |
|---|---|---|---|---|
| 1 | *Neurospora crassa* | 5141 | TrEMBL | euk |
| 1 | *Oryza sativa* | 4530 | TrEMBL | euk |
| 1 | *Ostreococcus tauri* | 70448 | Refprot | euk |
| 1 | *Paramecium tetraurelia* | 5888 | Refprot | euk |
| 1 | *Penicillium chrysogenum* | 5076 | TrEMBL | euk |
| 2 | *Piromyces finnis* | 1754191 | Refprot | euk |
| 1,2 | *Piromyces sp.* E2 | 73868 | TrEMBL | euk |
| 1 | *Saccharomyces cerevisiae* | 4932 | Refprot | euk |
| 3 | *Schizosaccharomyces cryophilus* | 866546 | TrEMBL | euk |
| NA | *Schizosaccharomyces japonicus* CBS5679 | NA | This study | euk |
| 3 | *Schizosaccharomyces japonicus* yFS275 | 402676 | TrEMBL | euk |
| 3 | *Schizosaccharomyces octosporus* | 4899 | TrEMBL | euk |
| 1 | *Schizosaccharomyces pombe* | 4896 | Refprot | euk |
| 1 | *Tetrahymena thermophila* | 5911 | Refprot | euk |
| 1 | *Ustilago maydis* | 5270 | Refprot | euk |

110

**Table S4: Observed compounds by GC-MS analysis. For mass spectra please refer to SI Dataset S06.**

All compounds detected by mass spectrometry in yeast biomass in this study, regardless of specific culture conditions, are listed in this table. The only exception is ergosta-5,7,22-trien-3β-ol, which was detected in a commercial ergosterol stock (see also Figure S2 of this SI Appendix datafile).

RRT = relative retention time; TMS = trimethyl-silyl

| Compound details | | | | GC-MS details | | | | Other | |
|---|---|---|---|---|---|---|---|---|---|
| Code | IUPAC Name | Common Name | Molecular weight [g/mol] | RRT cholestane | RRT ergosterol | RRT Ergosterol TMS ether | Characteristic ions and **Base peak** [*m/z*] | References | Comment |
| 1 | (6E,10E,14E,18E)-2,6,10,15,19,23-Hexamethyl-tetracosa-2,6,10,14,18,22-hexaene | Squalene | 410 | 0.94 | 0.73 | 0.71 | 410, 121, 95, 81, **69** | 6,7 | |
| *2* | 5α-Cholestane | Cholestane | 372 | 1.00 | 0.78 | 0.76 | 372, 357, 262, **217** | 6 | Reference material |
| *3* | 2,2-Dimethyl-3-((3E,7E,11E,15E)-3,7,12,16,20-pentamethylhenicosa-3,7,11,15,19-pentaenyl)-oxirane | Squalene epoxide | 426 | 1.07 | 0.83 | 0.81 | 426, 121, 93, **81**, 69 | 6 | |
| 4 | Hop-17(21)-ene | | 410 | 1.22 | 0.95 | 0.93 | 410, 395, **367**, 340, 231, 191, 189, 161, 135, 121 | 8 | |
| 5 | | | 428 | 1.28 | 0.99 | 0.97 | 428, 410, 395, 340, 243, **191**, 161, 135, 95 | | Structure possibly containing hydroxy group |
| 6 | Ergosta-5,7,22-trien-3β-ol | Ergosterol | 396 | 1.31 | 1.00 | 0.97 | 396, 378, **363**, 253 | | Reference material |
| 7 | Ergosta-5,7,22-trien-3β-ol TMS ether | Ergosterol TMS ether | 468 | 1.32 | 1.03 | 1.00 | 468, 378, **363**, 337 | | Through derivatization of **6** |
| 8 | | | 410 | 1.34 | 1.04 | 1.02 | 410, 395, **259**, 231, 189, 95 | | |
| 9 | | | 410 | 1.37 | 1.07 | 1.04 | 410, 395, 257, **243**, 231 | 9 | No reference data available |
| 10 | 4,4,14-Trimethylcholesta-8,24(28)-dien-3β-ol TMS ether | Lanosterol-TMS-ether | 498 | 1.43 | 1.09 | 1.08 | 498, 483, **393**, 241 | | Through derivatization of **11** |
| 11 | 4,4,14-Trimethylcholesta-8,24(28)-dien-3β-ol | Lanosterol | 426 | 1.44 | 1.10 | 1.10 | 426, 411, **393**, 241 | | |
| 12 | Hop-22(29)-ene | Diploptene | 410 | 1.45 | 1.12 | 1.09 | 410, 395, 367, 299, **191**, 189, 95 | 7,8 | Reference material |
| 13 | Hop-21(22)-ene | | 410 | 1.47 | 1.15 | 1.12 | 410, 395, 367, 341, 297, 231, **191**, 189, 161, 121 | 7 | |
| 14 | Hopan-22-ol | Diplopterol | 428 | 1.65 | 1.28 | 1.25 | 428, 395, 367, 341, 281, 207, **191**, 189, 149, 95 | 8 | |
| 15 | Hopan-22-ol TMS ether | Diplopterol TMS ether | 500 | 1.69 | 1.32 | 1.28 | 395, 340, 280, 191, 189, **131**, 95, 73 | | Through derivatization of **14** |

**Table S5. Measurements of glucose concentrations at various time points throughout an anaerobic growth study with *S. cerevisiae* strains.** An anaerobic pre-culture on synthetic medium with higher glucose (50 g $L^{-1}$ glucose) in the absence of sterols or UFAs was used to inoculate cultures in SMPD with 20 g $L^{-1}$ glucose, supplemented with Tween 80 and ergosterol (TE), Tween 80 only (T), ergosterol only (E) or none of these anaerobic growth factors (--). The first cultures of strains IMX2616 and IMX2629 supplemented with Tween 80 only (T1) were transferred to fresh medium of the same composition during the exponential phase (T2). b.d., below detection.

| Strain | Experiment | Sample timepoint (h) | Glucose concentration (mM) |
|---|---|---|---|
| **CEN.PK113-7D** | Anaerobic pre-culture | 18 | 67.48 ± 1.79 |
| | -- | 58 | 84.63 ± 0.25 |
| | T, mid sample | 23 | 77.32 ± 1.49 |
| | T, end sample | 58 | 35.54 ± 1.49 |
| | TE | 37 | b.d. |
| **IMX2616** *(sga1Δ::Sjshc1)* | Anaerobic pre-culture | 18 | 23.11 ± 0.35 |
| | -- | 58 | 97.69 ± 2.44 |
| | T1, mid sample | 23 | 17.42 ± 5.14 |
| | T1, end sample | 42 | 1.08 ± 1.02 |
| | T2 | 26 | 19.94 ± 0.63 |
| | TE | 37 | b.d. |
| **IMX2629** *(sga1Δ::Sjshc1, X-2Δ::Maths)* | Anaerobic pre-culture | 18 | 18. 97 ± 1.35 |
| | -- | 58 | b.d. |
| | T1, mid sample | 23 | 1.84 ± 0.94 |
| | T1, end sample | 42 | 0.09 ± 0.09 |
| | T2 | 26 | 17.42 ± 5.14 |
| | TE | 37 | b.d. |

### Supplementary Tables accompanying Materials & Methods

**Table S6: Relevant genotypes of *S. cerevisiae* strains used in this study**

| Strain name | Relevant genotype | Parent | Reference |
|---|---|---|---|
| CEN.PK113-7D | *MATα URA3 TRP1 LEU2 HIS3* | - | [10] |
| IMX2600 | *MATα URA3 TRP1 LEU2 HIS3*<br>*can1Δ::Cas9-natNT2* | CEN.PK113-7D | This study |
| IMX2616 | *MATα URA3 TRP1 LEU2 HIS3*<br>*can1Δ::Cas9-natNT2*<br>*sga1Δ::pTEF1-SjSHC-tCYC1* | IMX2600 | This study |
| IMX2629 | *MATα URA3 TRP1 LEU2 HIS3*<br>*can1Δ::Cas9-natNT2*<br>*sga1Δ::pTEF1-SjSHC-tCYC1*<br>*x2Δ::pTDH3-MaTHS-tADH1* | IMX2616 | This study |

**Table S7: Plasmids used in this study**

| Plasmid name | Relevant characteristics | Reference |
|---|---|---|
| p426TEF | 2μm AmpR *URA3 pTEF1-tCYC1* | [11] |
| p414-TEF1p-cas9-CYC | AmpR *TRP1 pTEF1-cas9-tCYC1* | [12] |
| pUG-natNT2 | AmpR *natNT2* | [13] |
| *pROS12* | 2μm AmpR *hphNT1* gRNA-*CAN1*.Y gRNA-*ADE2*.Y | [14] |
| pUD63 | 2μm AmpR *URA3 pTDH3-tADH1* | [15] |
| pUDR119 | 2μm AmpR *amdSYM* gRNA-*SGA1.Y* | [16] |
| pUDR538 | 2μm AmpR *hphNT1* gRNA-*X2.Y* | This study |
| pUD1150 | AmpR *MaTHS* | [ordered from GeneArt] |
| pUD1151 | AmpR *SjSHC* | [ordered from GeneArt] |
| pUDE1059 | 2μm AmpR *URA3 pTEF1-SjSHC-tCYC1* | This study |
| pUDE1060 | 2μm AmpR *URA3 pTDH3-MaTHS-tADH1* | This study |

**Table S8: Oligonucleotide primers used in this study**

| Primer # | Sequence |
|---|---|
| 2873 | TCAGACTTCTTAACTCCTGTAAAAACAAAAAAAAAAAAAAGGCATAGCAATAAGCTGGAGCTCATAGC TTC |
| 3093 | ACTATATGTGAAGGCATGGCTATGGCACGGCAGACATTCCGCCAGATCATCAATAGGCACCTTCGTAC GCTGCAGGTCGAC |
| 3903 | GCGAATTTCTTATGATTTATGATTTTTATTATTAAATAAG |
| 4653 | GTGCCTATTGATGATCTGGCGGAATGTCTGCCGTGCCATAGCCATGCCTTCACATATAGTCCGCAAAT TAAAGCCTTCGAG |
| 5542 | CTATGCTACAACATTCCAAAATTTGTCCCAAAAAGTCTTTGGTTCATGATCTTCCCATACGCATAGGC CACTAGTGGATCTG |
| 5921 | AAAACTTAGATTAGATTGCTATGCTTTCTTTCTAATGAGC |
| 6005 | GATCATTTATCTTTCACTGCGGAGAAG |
| 7298 | TTGTTCAATGGATGCGGTTC |
| 7376 | GGTCTAGGCCTGCATAATCG |
| 7377 | TGCGGCATCATGTCTACTTG |
| 7479 | GGACGTTCCGACATAGTATC |
| 9034 | TCACAGAGGGATCCCGTTACCCATCTATGCTGAAGATTTATCATACTATTCCTCCGCTCGGCGATCGC GTGTGGAAGAAC |
| 10546 | ATCCGTCGAAACTAAGTTCTG |
| 10547 | TCATGTAATTAGTTATGTCACGC |
| 10866 | TGCGCATGTTTCGGCGTTCGAAACTTCTCCGCAGTGAAAGATAAATGATCGGCGACTAGGAAGAGAG TAGGTTTTAGAGCTAGAAATAGCAAGTTAAAATAAG |
| 15002 | AATAAGTATATATATATTTGATGTAAATATCTAGGAAATACACTTGTGTATACTTCTCGCACCGGCC GCAAATTAAAGC |
| 15003 | TTTACAATATAGTGATAATCGTGGACTAGAGCAAGATTTCAAATAAGTAACAGCAGCAAAGCTGGAG CTCATAGCTTC |
| 15100 | GCTCATTAGAAAGAAAGCATAGCAATCTAATCTAAGTTTTATGAAAACTGATGGTAACAC |
| 15101 | GGAGGGCGTGAATGTAAGCGTGACATAACTAATTACATGATCAGAAGGCGAAAGAAACAT |
| 15102 | CTGCTGTTGCTGCTATGTTG |

| | |
|---|---|
| 15103 | TGGGTTGATAGCCTTTGG |
| 17519 | TTTTTTAGTTTTAAAACACCAGAACTTAGTTTCGACGGATATGACAAAAAATGCCAACTT |
| 17520 | CTTATTTAATAATAAAAATCATAAATCATAAGAAATTCGCTTAAGATGGGTAAGATTCTGGG |
| 17521 | GTCATAACTCAATTTGCCTATTTCTTACGGCTTCTCATAAAACGTCCCACACTATTCAGGCTTCCGGC TCCTATGTTGTG |
| 17522 | AACGACCACTTAGGTGAAGG |
| 17523 | GTACTTCAAGGCTGGATGAC |

For access to supplementary datasets, please contact the authors

**Dataset S01 (separate Excel file).**

Average values and mean deviation for triterpenoid- and TFA content of yeast biomass from duplicate aerobic and anaerobic growth experiments.

**Dataset S02 (separate Excel file).**

HMMER results using an oxidosqualene cyclase (OSC) from *Sch. pombe* (SpErg7; UniProt accession Q10231), a squalene-hopene cyclase (SHC) from *Acidocaldarius alicyclobacillus* (AaShc; P33247), and a squalene-tetrahymanol cyclase (STC) from *Tetrahymena thermophila* (TtThc1; Q24FB1), against eukaryotic and bacterial protein sequence databases.

**Dataset S03 (separate FASTA file).**

Amino acid sequences used to generate Figure 4. File provided in FASTA format.

**Dataset S04 (separate PHY file).**

Raw phylogenetic tree used to generate Figure 4. File provided in PHYLIP format.

**Dataset S05 (separate Excel file).**

HMMER E-values and corresponding sequence identifiers using CBS5679 either SjShc1 or SjErg7 as query against bacterial, and eukaryotic protein sequence databases.

**Dataset S05 (separate text file).**

Raw phylogenetic tree used to generate Figure 4. File provided in PHYLIP format.

**Dataset S06 (separate PDF file).**

Recorded mass spectra of compounds detected by GC-MS in yeast biomass in this study.
